# Metapopulations, the inflationary effect, and consequences for public health

**DOI:** 10.1101/2023.10.30.564450

**Authors:** Nicholas Kortessis, Gregory Glass, Andrew Gonzalez, Nick W. Ruktanonchai, Margaret W. Simon, Burton Singer, Robert D. Holt

**Author notes:** **Author emails**: NK; GG; AG; NWR; MWS; BS; RDH. **Author contributions**: NAK and RDH conceived on the theme of the manuscript. NAK wrote most of the initial draft, with original writing contributions from other authors. Editing of the initial draft was done collectively by all authors.

## Abstract

The metapopulation perspective is an important conceptual framework in ecology and evolutionary ecology. Metapopulations are spatially distributed populations linked by dispersal. Both metapopulation models and their community and ecosystem level analogues, metacommunity and meta-ecosystem models, tend to be more stable regionally than locally and display enhanced abundance because of the interplay of spatiotemporal heterogeneity and dispersal (an effect that has been called the “inflationary effect”). We highlight the essential role of spatiotemporal heterogeneity in metapopulation biology, sketch empirical demonstrations of the inflationary effect, and provide a mechanistic interpretation of how the inflationary effect arises and impacts population growth and abundance. We illustrate the effect with examples from the spread of infectious disease. Namely, failure to recognize the full possible effects of spatiotemporal heterogeneity likely enhanced the spread of COVID-19, a failure based on lack of understanding of emergent population processes at large scales which may hamper control and eradication of other infectious diseases. We finish by noting how the effects of spatiotemporal heterogeneity have implicitly played roles in the history of ecology, ranging across subdisciplines as diverse as natural enemy-victim dynamics, species coexistence, and conservation biology. Seriously confronting the complexity of spatiotemporal heterogeneity could push many of these subdisciplines forward.

## Introduction

The concept of a metapopulation – a set of spatially distributed populations linked by dispersal – is central to discourse in ecology, evolution, and conservation biology.

Metapopulation theory (Hanski and Gaggiotti, 2004)—and its community-level counterpart, metacommunity theory (Leibold, 2018)— provide a foundation for addressing important issues in ecology, including population persistence (Levins, 1969), species coexistence (Amarasekare, 2003; Hoopes et al., 2005), community assembly (Bergmann and Leveau, 2022; Leibold et al., 2022), and community stability (Loreau et al., 2003; Thompson et al., 2017). Metapopulation models have also helped in advancing our understanding of biogeographic patterns (Gonzalez et al., 1998) and evolutionary processes such as the ecology and evolution of range limits (Holt and Keitt, 2000), dispersal (Olivieri et al., 1995; Wang and Altermatt, 2019), and the interplay of selection and gene flow in adaptive evolution (Hanski et al., 2011; Ingvarsson, 2002). In applied contexts, metapopulations have been used to understand biocontrol (Levins, 1969), habitat loss, and habitat fragmentation (Ovaskainen and Hanski, 2004).

The origins of metapopulation theory come from observations by empirical ecologists such as Andrewartha and Birch (1954), Huffaker (1958) and den Boer (1968) on the often ephemeral and unstable nature of local populations. The formal theory, stemming back to the seminal paper of Levins (1969) for a single species, and later extended to interacting sets of species in metacommunities (e.g., Holt, 1997; Tilman et al., 1997), addressed the challenge of understanding the interplay of dispersal and environmental heterogeneities. The complex spatial and temporal structure of environments shapes the fitness of individuals and biology of species over both ecological and evolutionary time scales.

Like all valuable theory, metapopulation and metacommunity theory neglects many complexities of the world to make other complexities manageable. Metapopulation models simplify space into suitable habitat patches surrounded by non-habitat, akin to islands in an ocean. Early representations of metapopulations simplified population growth by representing any given patch as either containing a species (i.e., occupied) or not (i.e., unoccupied), a simplification that persists in many modeling efforts today (e.g., occupancy-based models). By ignoring details of spatial variation and the intricacies of local population growth, metapopulation models facilitate the study of population and community dynamics at large scales. Encompassing the full details of continuous space and population dynamics, even without spatial and temporal environmental heterogeneities, often leads to unwieldy mathematical problem (see e.g., Bolker and Pacala, 1997; Holmes et al., 1994). The simplifying assumptions of metapopulation models make them tractable, but they nonetheless retain some essential features of heterogeneities in space and time.

But not all. The simplicity of metapopulation models is, in some senses, deceptive. Our purpose here in effect is to ‘look under the hood’ of such models and bring out key but largely unrecognized assumptions crucial to their behavior. We will highlight how recognition of these assumptions, especially the nature of environmental variation, can explain emergent population properties at broad temporal scales – ignorance of which degraded the ability of public health authorities to manage and control the SARS-CoV-2 pandemic around the globe.

First, we reflect on the history of theory related to metapopulations and metacommunities and highlight the implicit role of spatiotemporal heterogeneity in their dynamics. Without spatiotemporal heterogeneity, many of the foundational lessons from classic metapopulation theory no longer hold. Then, we highlight how the presence of spatiotemporal heterogeneity influences multi-species counterparts to metapopulations—metacommunity models. We next explore a phenomenon that has received some attention in the theoretical literature but little empirical attention—namely the inflationary effect (a term coined in Gonzalez and Holt, 2002). The inflationary effect refers to the elevated growth rate (Kortessis et al., 2020) or population abundance (Gonzalez et al., 1998) that emerges in spatiotemporally heterogeneous landscapes coupled by dispersal, as compared to the same landscapes without temporal heterogeneity (Gonzalez and Holt, 2002). Studies of the inflationary effect attempt to deal more explicitly with heterogeneities in space and time and illustrate a phenomenon that we suggest is implicit in many metapopulation models. We discuss how awareness of the inflationary effect enriches our understanding of the spread of infectious disease, and how a lack of such awareness hampered the effective management of the pandemic caused by SARS-CoV-2. We finish by identifying future directions in ecology, evolution, and conservation biology that involve spatiotemporal heterogeneity and the inflationary effect. Our emphasis throughout is on conceptual issues and illustrative examples, not on mathematical details, although we provide key references in the history of the literature that provide such details for interested readers.

### Spatiotemporal heterogeneity

Metapopulation models embody a host of implicit assumptions about the nature of the environment. Amongst them, perhaps the most consequential is that spatiotemporal heterogeneity is influential for the dynamics of species. Spatiotemporal heterogeneity denotes how the temporal pattern of conditions varies as one moves across space. Spatial models alone cannot capture the effects of spatiotemporal heterogeneity. Nor can models with temporal heterogeneity, assuming a spatially well-mixed population. Spatiotemporal heterogeneity is a unique aspect of the environment revealed only when considering space and time together. Though the founders of metapopulation theory certainly recognized this dimension of theory (e.g., Hanski, 1998), an explicit discussion of this fundamental assumption is encountered infrequently in the literature, and so deserves special attention.

Before delving into models, it is first worth being specific about spatiotemporal heterogeneity: What is it, exactly? In essence, spatiotemporal heterogeneity occurs when patterns of conditions relevant to fitness over time differ across space (or spatial patterning that changes over time). Spatiotemporal heterogeneity can be expressed by a statistical decomposition of the total variation in some fitness-affecting factor, *F*(*x,t*), with both time *t* and location *x* coordinates (illustrated in Fig. 1a). The total variance in this factor, *σ*^2^ = Var(*F*), can be broken into three parts: *σ*^2^*_S_*, variability in space that is present after averaging over time, *σ*^2^*_T_*, variability in time that is present after averaging over space, and *σ*^2^*_ST_*, spatiotemporal variability (see Box 1 for a mathematical description and Chesson, 1985; Johnson and Hastings, 2023; Melbourne et al., 2007; Schreiber, 2023 for alternative, but conceptually similar definitions).

**Figure 1.**
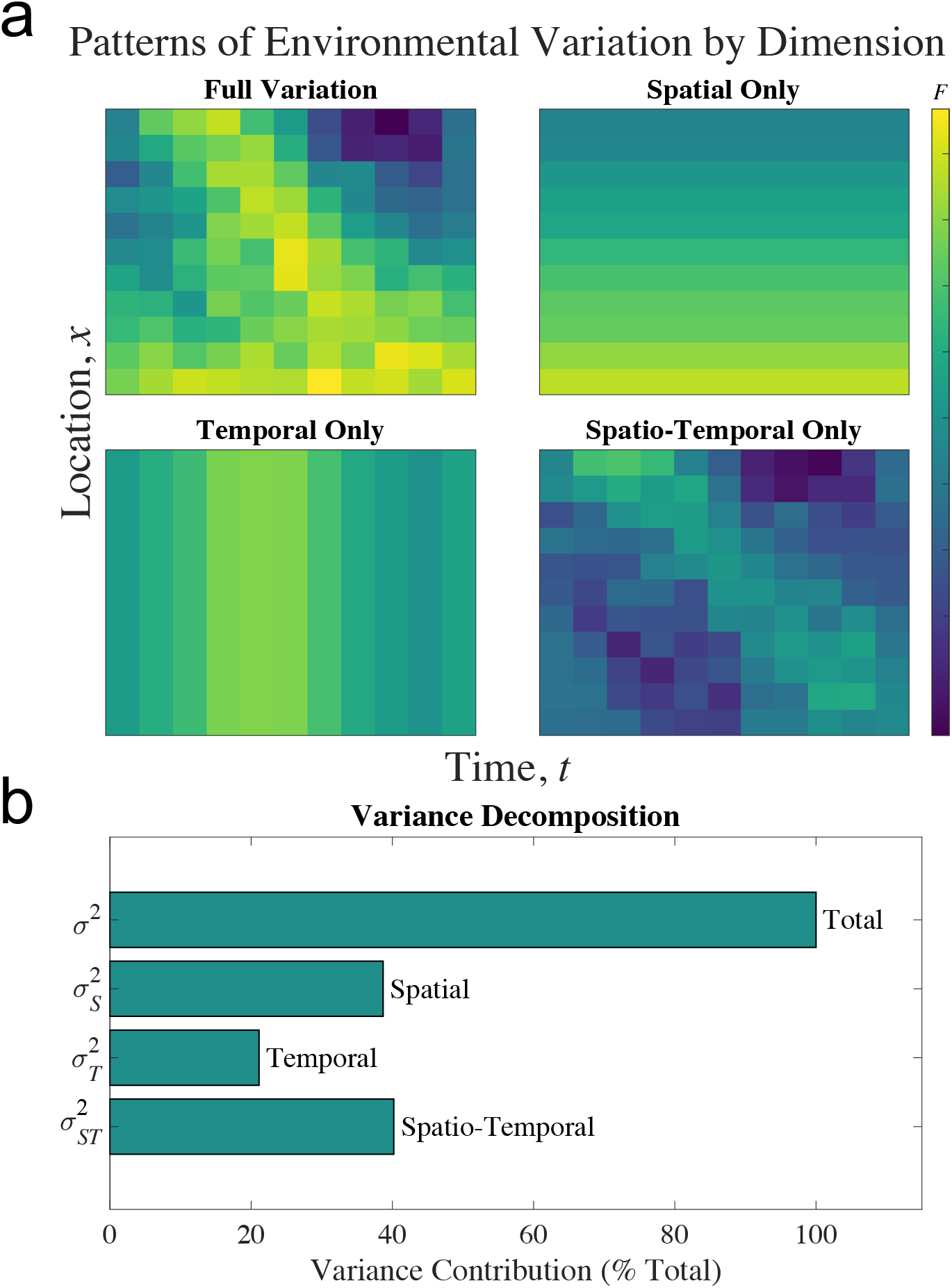
A hypothetical landscape with temporal and spatial variability in some fitness factor. (a) Spatial locations are aligned vertically and time points are aligned horizontally. Averaging values across time (i.e., horizontally) gives 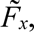 a perspective that illuminates average differences across locations. Averaging values across space (i.e., vertically) gives 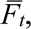 a perspective that illuminates average differences over time. The remaining variation is spatiotemporal and reflects differences in the pattern of variation across spatial locations. (b) The decomposition of the total variation across space and time in the fitness factor for the landscape in (a).

#### Box 1. Mathematical description of spatiotemporal heterogeneity

To illustrate how to quantify spatiotemporal heterogeneity, consider a landscape with locations indexed by *x* and times indexed by *t*. Let *F*(*x*,*t*) be a measure of some factor that influences the fitness or growth rate of individuals found in location *x* at time *t*.

A characterization of these environmental factors that does not include the effects of time is a *spatial-only* perspective. Consider the probability distribution for a fitness-factor in a location *x*, which we denote with the conditional variable *F*|*x*. A reasonable measure for the conditions in location *x* is 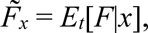 the time-average of the factor in location *x* (*E_t_*[⋅] being the expectation with respect to time; we use a tilde to indicate a time average). The variation present in space of these values, 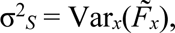 measures *pure spatial variation* (Var*_x_*(⋅) is the variance with respect to space).

A characterization of variation ignoring space is a *temporal-only* perspective. A reasonable measure of the conditions at time *t* across the entire landscape is 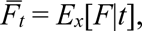 the spatial average conditions at time *t* (*E_x_*[⋅] being the expectation with respect to space; we use an overbar to indicate a spatial average). The variation present across time, 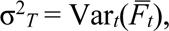 gauges *pure temporal variation* in the factor affecting population growth (Var*_t_*(⋅) is the variance with respect to time).

With these definitions of pure spatial and pure temporal variance in hand, we can write the total variation in the factor affecting population growth as

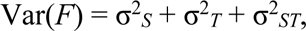

where σ^2^*_ST_* measures *spatiotemporal variation*. We can define this spatiotemporal variation as (see supplementary material S1 for derivation)

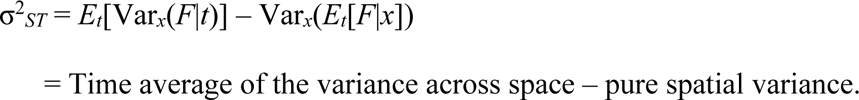

The quantity σ^2^*_ST_* represents the amount of excess variation that occurs when we look at an ensemble of spatial variation across time rather than when averaged over time. An equivalent (and mathematically identical) representation quantifies the excess variation in time that occurs when we look at an ensemble across space over a simple average across space (see supplementary material S1 for details).

The term σ^2^*_ST_* in Box 1 quantifying heterogeneity arising jointly across space and time is our focus. We will argue that this is not a merely ‘residual’, but has important consequences.

Spatiotemporal heterogeneity cannot be explained by spatial-only and temporal-only perspectives of the world, separately or added up (Fig. 1a-b). Figure 1a visualizes the decomposition. In a temporal-only perspective, conditions vary over time and are the same across space. In a spatial-only perspective, conditions differ across space but are maintained over time. Spatiotemporal variation is additional variation on top of these two sources and may constitute anywhere from 0% of the total variability to 100%.

Spatiotemporal heterogeneity can be generated in many ways. One fundamental source of spatiotemporal heterogeneity is interactions between physical variation in abiotic factors across space and time. For example, temperature fluctuations differ among land slopes with different aspects; daily and seasonal fluctuations in temperature can be greater on south-facing slopes in the northern hemisphere, and on north-facing slopes in the southern hemisphere. In aquatic metacommunities in landscapes with many lakes and ponds differing in area and depth, larger and deeper water bodies are buffered from fluctuations in physical factors such as temperature, compared to smaller and shallower water bodies.

In addition to physical factors, biotic factors can create strong spatiotemporal heterogeneity. Any species interacting with a focal species—biotic resources, natural enemies, competitors, and mutualists—can generate variation in that focal species’ fitness. Fluctuations in the abundance or activity levels of an interacting species can potentially lead to spatiotemporal heterogeneity in the fitness of a focal species if such fluctuations play out differently in different locations. Spatiotemporally varying abundances of species are often said to have spatially *asynchronous* dynamics (Shoemaker et al., 2022). Asynchronous population dynamics are a form of spatiotemporal heterogeneity because the dynamical patterns of abundance fluctuations differ across space (although perfectly synchronous dynamics can exhibit spatiotemporal heterogeneity if the magnitude of synchronized fluctuations differ across locations).

Asynchronous dynamics can arise even in homogeneous physical environments, due to complex processes involving dispersal, chance events, and interactions with other species (Gouhier et al., 2010; Keeling et al., 2000; Levin, 1974; Vasseur and Fox, 2009). For example, chaotic dynamics could be a potent source of asynchrony even in physically homogeneous landscapes because slight differences in initial conditions lead to large deviations in resulting population trajectories among locations (Hassell et al., 1991; Holt, 1993). Regardless of the source of asynchrony, asynchronous dynamics of one species (e.g., a generalist predator) can drive spatiotemporal heterogeneity in the fitness of other species with which it interacts (e.g., one species of that predator’s prey).

### Metapopulations

One of the first approaches for dealing with spatiotemporal heterogeneity was to envision a metapopulation, a set of spatially discrete local populations prone to extinction and connected by dispersal. Metapopulations in the most general sense of the word are “populations of populations” (Hanski, 1999; Levins, 1970). They are populations defined on a scale that encompass sub-populations, the dynamics of which are determined by individuals present locally but also by individuals that leave (emigrate) and join (immigrate).

It is worth distinguishing distinct types of metapopulation models. The term metapopulation sometimes refers to models based solely on occupancy, and sometimes to models of patchily distributed populations where the abundances across locations are modeled. We will use the term “strict metapopulations”, to refer to models whose dynamics are characterized in terms of the qualitative state of the population (i.e., occupancy) and where recurrent local extinctions and colonizations determine dynamics (Hanski, 1998). We will use the term “patch models” to describe models in which dynamics are characterized by quantitative measures of population states (e.g., local density) and whose dynamics depend explicitly on birth, death, emigration, and immigration rates. Both models may include spatiotemporal variation, but its inclusion in patch models is often explicit whereas it is implicit in strict metapopulation models.

The distinction between “strict metapopulations” and “patch models” is to a degree an arbitrary one based on choices used to describe local populations. All real populations are spatially distributed with dynamics that could be described by abundance or by a simpler metric of abundance: presence (abundance greater than zero) or absence (abundance equal to zero). Indeed, any patch model could be rewritten as a strict metapopulation model, and there are many types of local population dynamics that are consistent with any given strict metapopulation model. They are, put simply, complementary modeling frameworks, each with their strengths and limitations.

### Simple models of spatiotemporal heterogeneity: spatial occupancy models

Levins (1969) introduced the concept of a metapopulation to deal precisely with the complexities generated by modeling spatiotemporal heterogeneity^1^. (The term “metapopulation” seems to have its first usage in Levins, 1970). He recognized that conditions differing across space and time could cause populations sizes to fluctuate wildly, oftentimes causing extinctions of local populations. But sites do not remain empty because reproductively viable individuals from nearby, extant populations can immigrate.

The localization of extinctions and the subsequent potential for recolonization have foundational roles in strict metapopulations. Classical, strict metapopulation models are based on the following canonical form:

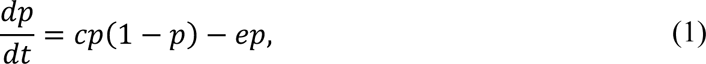

where *p* is the proportion of patches on the landscape currently occupied, *e* is the rate at which populations go locally extinct, and *cp* is the rate at which unoccupied patches again harbor a viable population (*c* is the colonization rate per occupied patch). Such a model describes landscapes with many patches whose extinction events are independent across patches, and where recolonization events do not depend on the spatial position of empty patches (Hanski, 1998, 1997). Model (1) predicts that if the extinction rate is lower than the colonization rate (i.e., *e* < *c*), the fraction of occupied patches grows logistically to an equilibrium of *p*\* = 1 – *e*/*c*. Even though local extinction is inevitable everywhere, frequent recolonizations allow for metapopulation persistence.

A crucial assumption is that local extinctions are not strongly synchronized across space; otherwise, recently extinct populations would have no sources for recolonization. This point has been made before (Chesson, 1981; Hanski, 1998; Hoopes et al., 2005), but is rarely a central point of discussion of metapopulations and so can be easily missed (although see Chesson, 1985; Harrison and Quinn, 1989; Petchey et al., 1997). The assumption in Levins’ model of no strong synchronization thus reflects an implicit assumption of the presence of spatiotemporal heterogeneity if environmental factors (rather than say pure demographic stochasticity) drive those extinctions. Conditions in any one local population, at any one time, are not identical to all other local populations during that time.

To illustrate this subtle assumption and its consequences for metapopulation dynamics, consider a finite patch version of model (1). The model has *n* patches where we track occupancy in each patch. Let *O_i_*(*t*) be the occupancy in patch *i* at time *t* (*O_i_* = 1 if occupied and *O_i_* = 0 if not), and let *P*(*t*) = ∑*_i_O_i_*(*t*)/*n* be the fraction of patches occupied at time *t*. To simulate the model, we consider how occupancy changes in a small unit of time, Δ*t*. In each unit of time, each occupied patch goes extinct with probability 1 – exp(–*e*Δ*t*), and each unoccupied patch becomes occupied with probability 1 – exp(–*cP*(*t*)Δ*t*). As the time steps get small, extinction occur at rate *e* per occupied patch, and colonization occurs at rate –*cP*(*t*) per unoccupied patch, just as in model (1) (see supplementary material S2 for model details.)

Spatiotemporal variability can be controlled with the *joint probability* of extinctions and colonizations across multiple patches each time step. If we assume extinction events in each patch are independent in space and time, we recover a finite-patch version of equation (1). In that case, any pattern of extinction-causing factors will have different patterns over time among patches (see supplementary figure S1), an example of strong spatiotemporal heterogeneity. If instead we assume extinction drivers in one patch correlate with drivers in other patches, patterns of extinction over time would share at least a degree of similarity across patches, i.e., there is decidedly less spatiotemporal heterogeneity. We can tune the level of spatiotemporal heterogeneity with *ρ_E_*, the correlation between any two patches in environmental conditions affecting extinction (for simplicity, we assume this correlation applies to all pairs of patches). Since all patches are otherwise the same, there is no spatial heterogeneity (*σ*^2^*_S_* = 0). When *ρ_E_* = 0, extinction events are independent across patches and all variation is spatiotemporal (*σ*^2^*_ST_* > 0; *σ*^2^*_T_* = 0). When *ρ_E_* = 1, extinction in one patch implies extinction in all, there is no spatiotemporal variation (*σ*^2^*_ST_* = 0). In fact, in that case, all variation is temporal (*σ*^2^ = *σ_T_*^2^).

With independent extinctions (i.e., *ρ_E_* = 0), the classical, strict metapopulation assumption is recovered, and the population reaches a relatively stable patch occupancy (Fig. 2, green line), even in a finite patch model with relatively few patches (*n* = 50). By breaking the assumption of independent extinction events across locations (i.e., reducing spatiotemporal heterogeneity), stability of the metapopulation weakens (Figure 2, green line). Occupancy exhibits large fluctuations because local extinctions, when they occur, tend to occur over a large spatial domain with many local populations impacted simultaneously. Even highly occupied metapopulations are vulnerable to drastic drops in occupancy because of highly synchronized extinction events.

**Figure 2.**
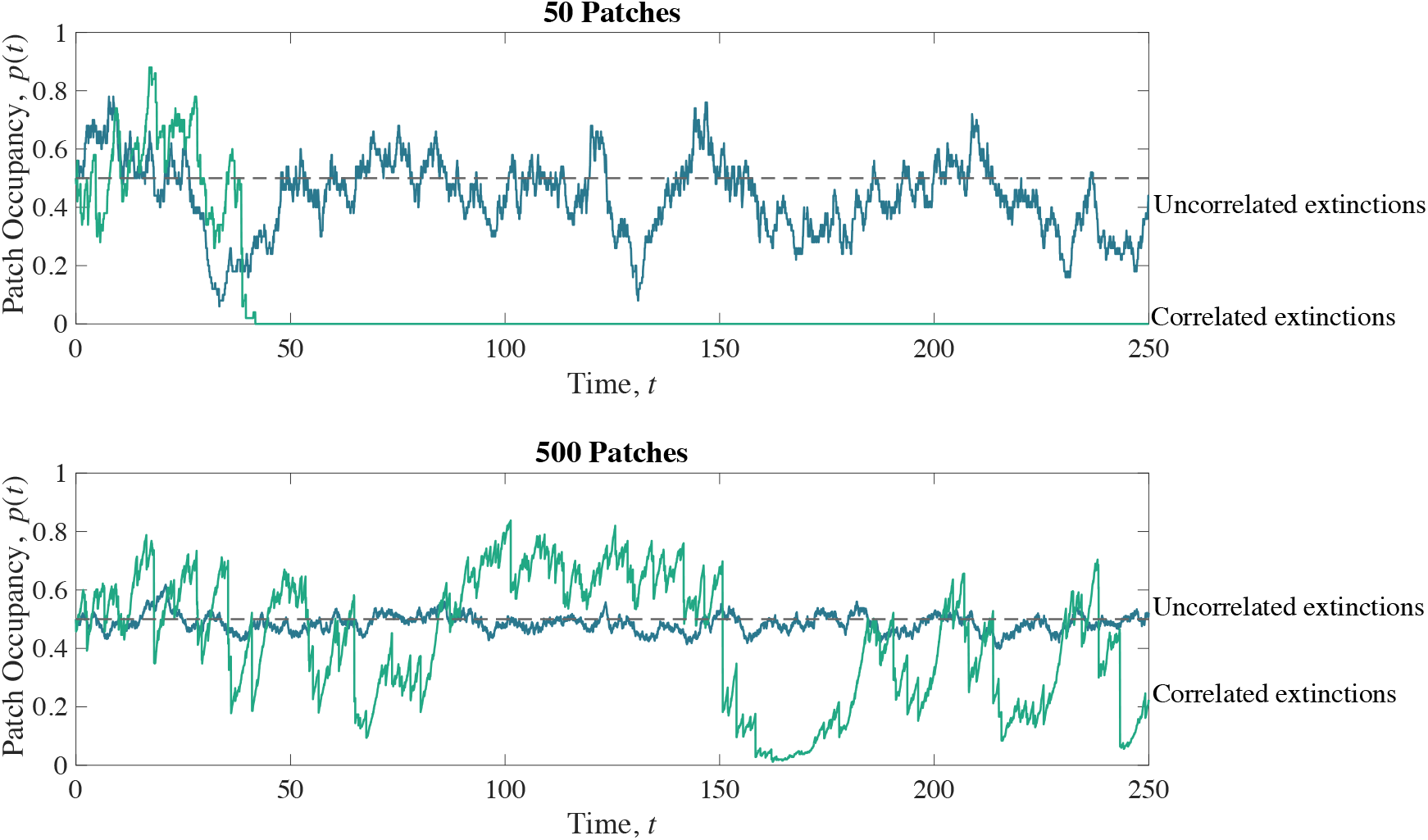
Finite patch version of Levins’ metapopulation model under different assumptions about asynchrony of local patch extinctions. Dynamics under the conventional assumption that environments influencing extinction are uncorrelated, i.e., asynchronous, with *ρ_E_* = 0, are given by the blue lines. Dynamics allowing for correlated extinctions, with *ρ_E_* = 0.75, are shown by the green lines. Simulation details found in supplement. For all dynamics, *e* = 0.25, and *c* = 0.5 such that the infinite patch model predicts equilibrium occupancy of *p*\* = 1 – *e*/*c* = 0.5 (dashed lines).

The large fluctuations caused by eroding spatiotemporal heterogeneity greatly increase the likelihood of metapopulation extinction. For example, a metapopulation of *n* = 25 patches with independent extinction events (i.e., strong spatiotemporal heterogeneity) persists an average of ∼300,000 units of time (Fig. 3a, dark purple points). Removing this implied spatiotemporal heterogeneity by increasing the correlation among patch extinction events reduces persistence times by several orders of magnitude (Fig. 3a), despite little change in average occupancy (Fig. 3b). This simple model shows that the well-known stability implied by metapopulation models results from the implicit assumption of strong spatiotemporal heterogeneity among patches.

**Figure 3.**
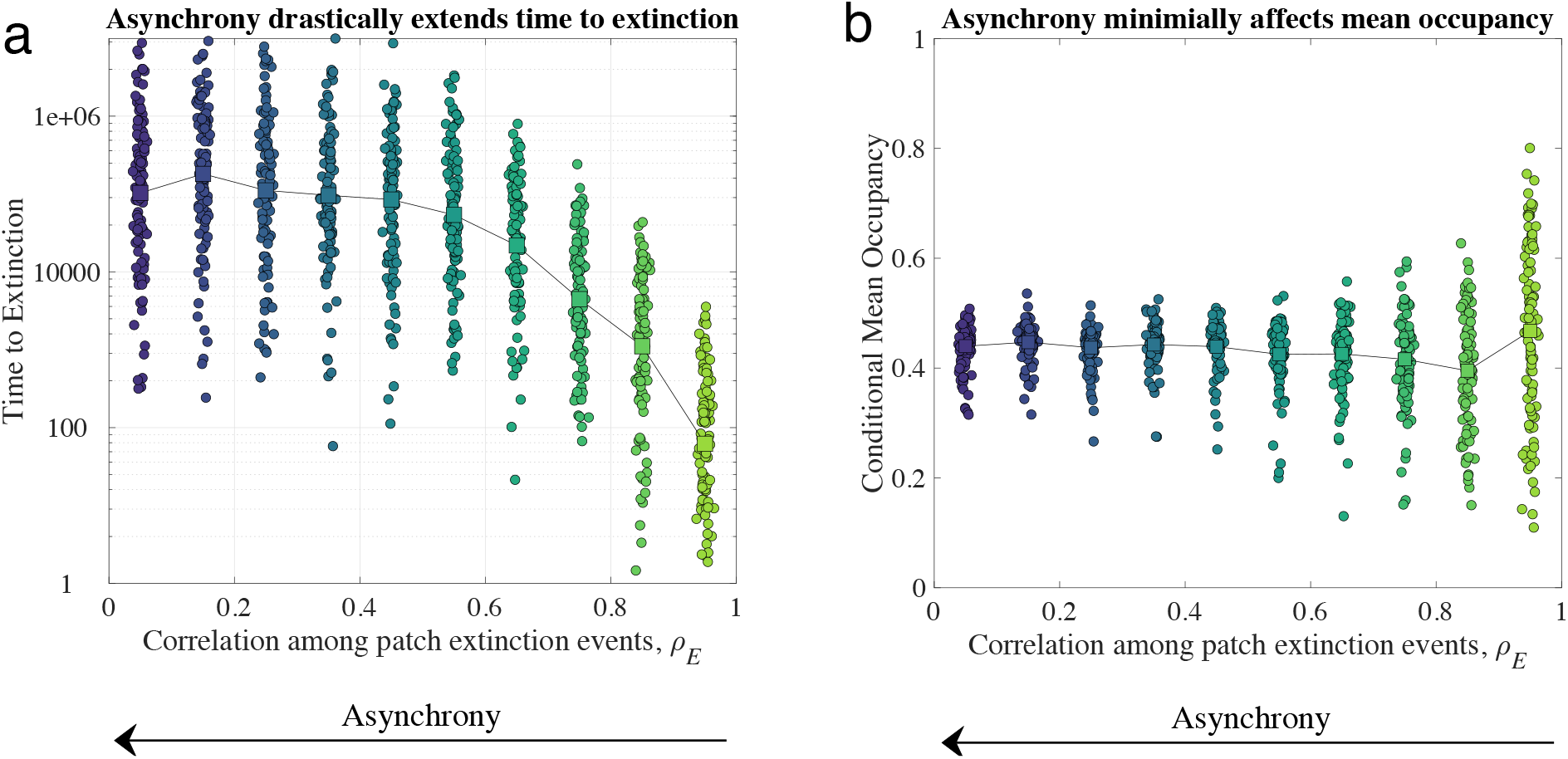
Effect of correlated extinction events on metapopulation stability. (a) Metapopulation extinction times for replicate simulations, and (b) corresponding conditional mean occupancy. Squares connected by lines are the geometric mean extinction times in (a) and mean conditional mean occupancy in (b) over all replicate simulations. The parameter *ρ_E_* refers to the correlation coefficient for all pairwise latent variables describing environmental variables determining extinction in patches (details in the supplementary material S2). All simulations had 25 patches, 13 of which were initially occupied. There are 100 replicate simulations for each *ρ_E_* value. Other parameters: *e* = 0.25, and *c* = 0.5.

### Metacommunities: spatiotemporal heterogeneity in species interactions

A natural extension of metapopulation models is to consider other species. Patches now represent local areas harboring communities connected by dispersal, the collection of which is a metacommunity (a term first used by Wilson, 1992). Species’ colonization and extinction rates in metacommunities reflect some combination of the physical environment, interspecies interactions, and demography. We distinguish “strict metacommunities”, whose dynamics are described in terms of extinction and colonization rates (i.e., occupancies, hence co-occupancies) (see e.g., Hastings, 1980), and community patch models, whose dynamics are described in terms of birth, death, immigration, and emigration rates of each constituent species within each patch (see e.g., Holt and Chesson, 2016). Spatiotemporal heterogeneity in patch models—a rich and growing area of community ecology (Driscoll and Lindenmayer, 2009; Leibold, 2018; Leibold et al., 2022; Lu, 2021)—is considered below. Here we focus on strict metacommunities.

To illustrate issues that pertain more broadly to more complex multispecies models, we use a simple model for a core component of any metacommunity – a specialist predator interacting with its required prey species. A specialist predator-prey strict metacommunity model is

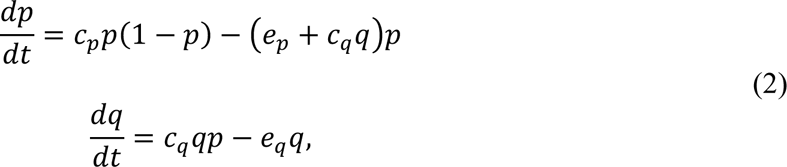

where *p* is the fraction of habitat patches occupied by prey alone, *q* is the fraction occupied by both predators and prey, *c_i_* is the colonization rate of patches of type *i* (*i* = *p,q*), and *e_i_* is the extinction rate of patches of type *i* (e.g., the prey alone, or it together with the predator).

Models such as (2) follow the same simple rules as a strict metapopulation model except that now extinction and colonization rates depend on other species. Patches occupied by prey alone change state independently of the predator (e.g., due to disturbance) with an extinction rate of *e_p_* and because they are colonized by the predator at a rate proportional to the availability of prey-only patches and the prevalence of predators (*c_q_pq*). We assume the predator greatly limits prey numbers in co-occupied patches, so that patches occupied by the predator do not provide any prey colonists to empty patches. The above model assumes that predators only go extinct when their prey go extinct (although more elaborate models that relax these assumptions to investigate other ways predators might influence prey extinction follow the same qualitative rules; see Holt, 1997).

Metacommunity models often exhibit stability despite locally unstable interactions between species in any patch (Gravel et al., 2016; Levin, 1974). Strong predator-prey dynamics typically generate large-amplitude fluctuations in spatially and temporally homogeneous environments (May, 1972). It is often difficult to sustain such dynamics experimentally (but see Blasius et al. 2020 for a recent laboratory success). But metacommunity models such as (2) exhibit stability despite such unstable local dynamics, where predators, once arriving at a patch, eventually drive prey extinct locally (Gouhier et al., 2010; Hastings, 1977).

The stability exhibited in metacommunity models again reflects spatiotemporal heterogeneity. To understand this, consider the predator first. Given sufficient availability of prey patches to colonize, predator stability is ensured by the fact that extinctions occur independently across locations, just as in metapopulation model (1). And a stable, suitable prey source is ensured by the same argument, but including the fact that predators also show some independence across patches in their colonization of prey-alone patches. Stated succinctly, predator effects are spatiotemporally heterogeneous. Synchronous predator colonization or extinction events would lead to large-scale oscillations comparable to those of Figure 2, with high likelihood of extinction in realistic models.

For other metacommunity models including, for instance, infectious disease and competitive interactions (discussed below), stability at the regional scale is also observed, and relies on the implicit assumption that extinction and colonization events are independent across patches, as in our stochastic metapopulation model (Figs. 2,3). Independence across patches requires spatiotemporal heterogeneity be present and strong. One perspective about strict metacommunity models, we suggest, is that some of the most interesting features about the maintenance of diversity in metacommunities emerge from spatiotemporal heterogeneity.

### Digging deeper: beyond models of occupancy

The stabilizing effects of spatiotemporal heterogeneity can be difficult to recognize in strict metapopulation models. Such effects become clearer in patch models, where population dynamics within patches are explicit in terms of births, deaths, emigration, and immigration. Rather than comprehensively review the large and diffuse literature related to this theme, we here follow one line of thought about the emergent effects of spatiotemporal variation.

Holt, Gonzalez, and other collaborators began exploring the consequences of spatiotemporal heterogeneity in patch models in a series of papers beginning in the early 2000s. In a simple but illuminating example, Gonzalez and Holt (2002) considered a single sink patch supported by constant immigration from a source patch. It had long been recognized that populations may exist in sinks given constant immigration from a source, and that the sink population could be substantial given moderate rates of population decline (Holt, 1985; Pulliam, 1988; Runge et al., 2006; Van Horne, 1983). Gonzalez and Holt (2002) expanded on this theme by envisioning that local sink conditions were sinks on average over time, but not all the time. On average, conditions in the sink habitat were too poor to sustain a population without immigration, but transient phases of locally favorable conditions might allow a population to temporarily grow. Without immigration, the population may fluctuate and temporarily increase, but it eventually, given its sink namesake, wanes away and goes locally extinct.

However, immigration prevents extinction during bad times, buffering declines to low numbers. Immigration then allows the persistent sink population to capitalize on good times, boosting time-average population sizes in the sink (sometimes by orders of magnitude), a phenomenon they termed “the inflationary effect”. Gonzalez and Holt (2002) demonstrated this effect mathematically and used a microcosm experiment with source-sink populations of the protozoan *Paramecium tetraurelia*, buffeted by temporally varying thermal conditions, to rigorously test the phenomenon.

The inflationary effect of spatiotemporal variation on abundance works as follows and is illustrated in Fig. 4. Conditions are sometimes poor for growth, and sometimes favorable for growth. Good conditions cause large upswings in population size, particularly if growth rates are positively autocorrelated, permitting runs of good conditions. Poor conditions decrease population size, but rather than the population crashing (and erasing any increase in abundance), persistent immigration into the sink maintains a minimum growth rate, keeping the population from dipping too low. Thus, the effects of favorable conditions outweigh the effects of unfavorable conditions, even if good and bad times are equally frequent. In essence, immigration maintains a minimum local growth rate that, when paired with temporarily favorable conditions for growth, greatly increases sink population size.

**Figure 4.**
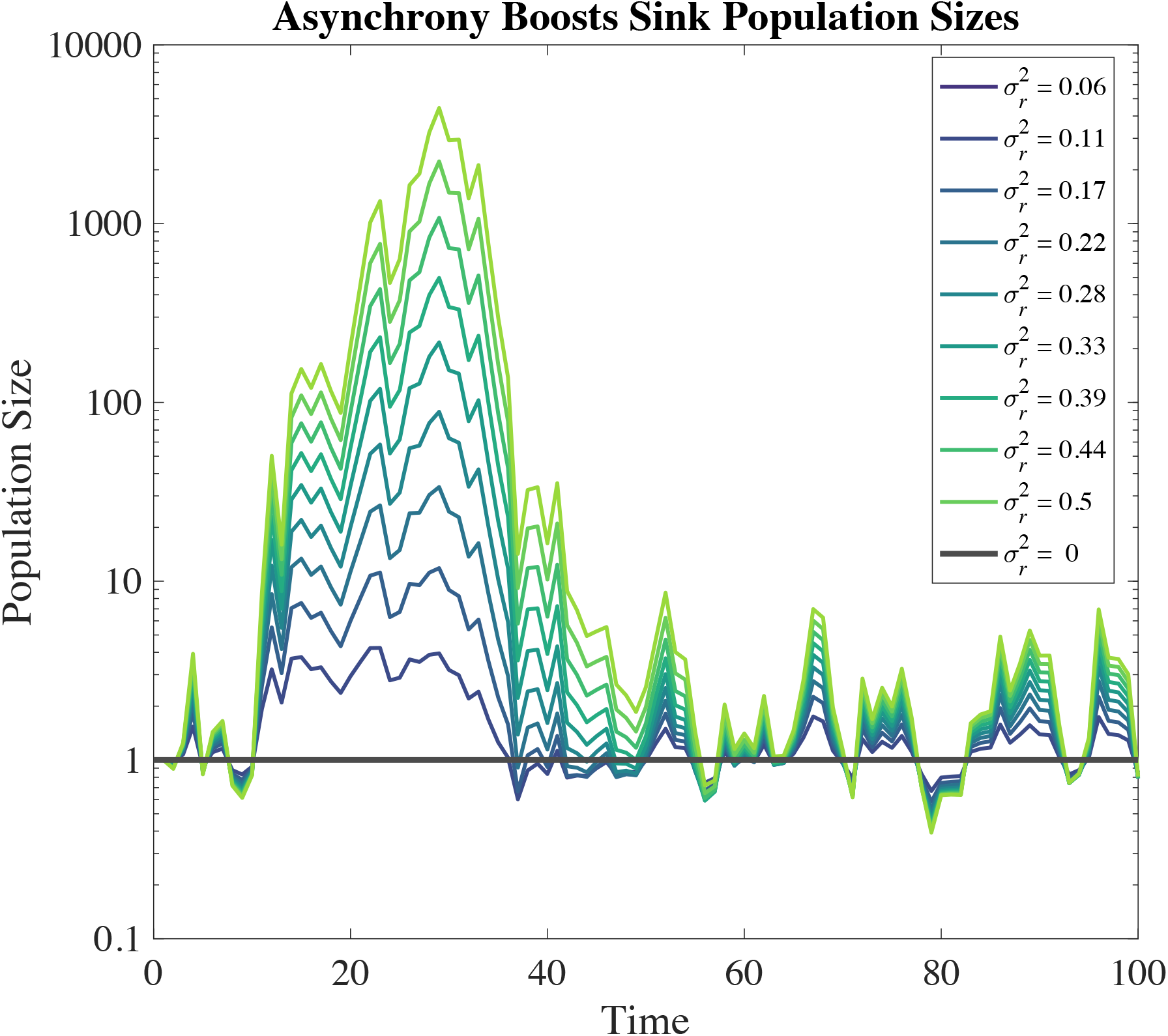
Sink population dynamics with fluctuating growth rate and a constant immigration. Dynamics follow *N*(*t*+1) = *N*(*t*)e*^r^*^(*t*)^ + *I*, where *r*(*t*) is the per-capita growth rate without immigration, and *I* the density of immigrants. Here 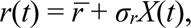 where *X*(*t*) is follows a standard normal distribution. We chose 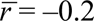 and *I* = 1 – *e*^-0.2^ ≈ 0.18 such that the equilibrium density of the sink in a constant environment is 1 (heavy black line). All cases start with *N*(*t* = 0) = 1 and have identical sequences of *X*(*t*). Further details in supplementary material S3.

This increase in sink population size – an inflationary effect of abundance – is fundamentally about spatiotemporal heterogeneity. The measure of spatiotemporal variance from Box 1 applied to the model first investigated in Gonzalez and Holt (2002) reveals that *σ*^2^*_ST_* = *σ_r_*^2^/4, where *σ_r_*^2^ is the variance in the sink growth rates (see supplementary material S3). Hence, the inflationary effect of population size in Fig. 4 is present when *σ*^2^*_ST_* > 0 and absent when *σ*^2^*_ST_* = 0 (black line in Fig. 4).

Further modeling studies demonstrated that the inflationary effect arises in many other, more complex and biologically realistic models, and may show up in the metapopulation growth rate as well as time-averaged abundance. The effect occurs in source-sink models with explicit resource dynamics, density-dependence, demographic stochasticity, and competition between species (Holt et al., 2003). Somewhat remarkably, it also applies to multiple sinks linked by dispersal (so there is no persistent source habitat anywhere in the landscape at all) and can be sufficiently strong in some cases to ensure the persistence of a regional landscape comprised entirely of stochastic sinks (Roy et al., 2005).

This last point is worth reiterating. The inflationary effect can allow for the regional persistence of populations living in a landscape of sinks. That is to say, in any given location on the landscape, a population isolated from immigration cannot persist. But if some individuals there disperse (not too many) to neighboring locations, such reshuffling of individuals can be sufficient to allow for all populations to persist in perpetuity. This boost in regional growth depends on spatiotemporal variation and asynchrony in growth conditions among patches, and it is a specific case of paradoxical persistence of mixed dynamics systems with strong heterogeneity (see the excellent review of such systems in Williams and Hastings 2011).

A microcosm study (Matthews and Gonzalez, 2007) prompted by this result convincingly demonstrated that asynchrony could permit metapopulation persistence, even for just two patches coupled by dispersal. Matthews and Gonzalez observed that two local populations of *Paramecium aurelia* would quickly go extinct in both patches for treatments with synchronous temperature fluctuations across patches, but populations persisted in treatments with asynchronous temperature fluctuations across the two patches. They also showed that positive autocorrelation boosted the inflationary effect on sink population sizes, a result that was extended to host-parasite dynamics (Duncan et al., 2013) in metapopulations of *Paramecium caudatum* and the bacterial parasite *Holospora undulata*. The key finding is that autocorrelated and asynchronous variation in temperature allowed infected host populations to maintain sizes equivalent to uninfected populations, despite negative demographic impacts on infected individuals.

One might reasonably wonder whether such results are limited to small parameter ranges with highly specific conditions, in which case the inflationary effect would merely be a theoretical curiosity of scant real-world importance. Recent studies however have investigated quite general models of population growth in discrete and continuous-time and find the inflationary effect to be present over broad ranges of parameter space (Benaïm et al., 2023; Katriel, 2022; Schreiber, 2010). Moreover, other studies have shown that the rates of dispersal consistent with the inflationary effect are adaptive in both single species (Schreiber, 2012) and multispecies models (Schreiber et al., 2023). These studies demonstrate a very general pattern: spatiotemporal heterogeneity in growth rates can robustly allow for persistence of species and increase their abundance. This should not come as too much of a surprise, as the lesson from theories of metapopulations is that the metapopulation can persist despite no individual patch persisting on its own, without the others. The inflationary effect is about just such persistence, but the details of these models demonstrate the mechanism by which spatiotemporal heterogeneity can be beneficial for population persistence, as we demonstrate below.

### What is “the inflationary effect”, and how does it work?

The clearest requirements for the inflationary effect were illustrated by Roy et al. (2005) for abundance and Schreiber (2010) for growth rate, which we summarize here as they provide a pathway to understand its mechanistic underpinnings. In both cases, the inflationary effect can be thought of as the effect of spatiotemporal heterogeneity inflating some aspect of population growth at the scale of the metapopulation, such that it is much larger than some simple average of local conditions.

Prior studies illustrate that the inflationary effect needs (1) some (but not too much) movement among patches, (2) variation in local growth rates over time, (3) a less than perfect spatial correlation in growth rates, and (4) temporal autocorrelation in growth rates. Temporal variation and low spatial correlation in growth rates together are signatures of spatiotemporal heterogeneity. The combination of movement and temporal autocorrelation together allows individuals in the metapopulation sufficient time to disperse across space so as to experience spatiotemporal heterogeneity. Without movement, the effective landscape of an individual is a single patch, not a metapopulation, despite the landscape having multiple patches. Without temporal autocorrelations, environments shift before populations can build up in transiently favorable locations. As populations grow, more individuals emigrate (in an absolute, not per-capita sense), even if the locations into which they immigrate are currently poor for growth. Given spatiotemporal heterogeneity, favorable conditions in one patch imply that emigrants from there are likely to immigrate to a temporary sink, boosting its numbers. Temporary sinks disproportionately benefit from immigration because it maintains minimum population sizes, which enhances population (not per-capita) growth rate when the environment eventually turns, as illustrated in the sink model with constant immigration above (Figure 4, see supplementary material S3 for model details).

This qualitative description can be made more precise with a mathematical description of the effect, which reveals how the mechanism operates on growth rates. Others have provided a mathematical description for discrete-time (Johnson and Hastings, 2023; Schreiber, 2023, 2010), but we present one for continuous-time where the math is (slightly) less painful. Consider a set of *n* connected patches where *N_i_* is the local density in a patch *i*, *r_i_*(*t*) is the local per capita growth rate, and *m_ij_* be the patch-specific per capita movement rates from patch *j* to *i*. The dynamics in patch *i* can be written as

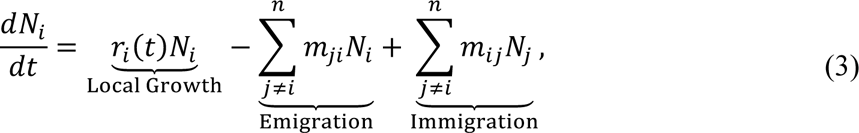

which allows for time- and location-dependent growth and movement rates.

To understand the role of spatiotemporal heterogeneity, we could ask how different population growth would be in the absence of spatiotemporal heterogeneity. Hence, a reference scenario in which spatial and temporal variation’s effects are present, but the spatiotemporal effects are not, is required. We use the scenario where the average growth rates across time describe local growth in each patch (using the notation in Box 1, 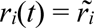 for all *t*). This scenario encapsulates the effects of temporal variability alone with the average growth rate and the effects of spatial variability by allowing each population to have different average growth rates with dispersal. This population reaches a stable spatial distribution, which is analogous to the stable age distribution in stage-structured demographic models. Let 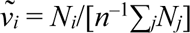 be the stable relative density in patch *i* in the reference scenario. The growth rate in this reference scenario can be found with the dominant eigenvalue of (3) (see supplementary material S4).

The amount to which the actual metapopulation growth rate deviates from this reference scenario without spatiotemporal heterogeneity is

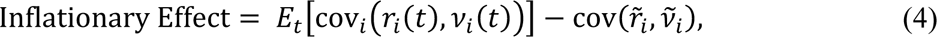

where *ν_i_*(*t*) = *N_i_*(*t*)/[*n*^-1^∑*_j_N_j_*(*t*)] is the relative density in location *i* at time *t* in the full model with spatio-tempoal heterogeneity (derivation in supplementary material S4).

Equation (4) measures the average association over time between the density of individuals across space and local growing conditions and how it compares to the association between average conditions. As such, it measures the resulting ability of populations to “track” good locations that shift in time and space in reference to the ability to find and persist in good locations on average. Expression (4) depends on a time-average of a covariance, which may be either positive or negative, and may lie above or below the covariance for the reference scenario, implying that the inflationary effect could be positive, zero, or negative. At any given time, the covariance is positive when individuals are concentrated in better than average spatial locations and is negative if instead individuals in the metapopulation are concentrated in poorer than average spatial locations. When population density is random with respect to local patch growth rates, the covariance is zero. Negative covariances can arise, for instance, under directed passive dispersal out of sources into sinks, as demonstrated by Keddy’s (1982) study on the sea rocket in Nova Scotia dunes, where seed dispersal is driven by winds from the ocean across the dunes.

Prior studies studying the inflationary effect have focused on situations where equation (4) is positive, meaning that a randomly chosen individual in time and space experiences conditions better than the average spatial conditions in time. These studies find that the average individual experiences (somewhere) conditions in space that are (temporarily) favorable to growth. These opportunities may not be present consistently, in any time or place, but are present ephemerally. Equation (4) measures how dispersal capabilities of species interact with the spatiotemporal structure of the environment. (Deriving explicit expressions for this quantity is analytically challenging, however some approximations are possible; Schreiber, 2023.)

Studies on the inflationary effect typically assume passive forms of non-directional dispersal, which leads the average individual to experience better than average conditions, even without dispersal adaptively directed towards such locations. A positive covariance between local growth rate and local relative abundance arises simply because populations tend to grow more where conditions are temporarily good. Adaptive tracking of the environment will certainly enhance the effect of spatiotemporal variation in fitness on population growth. For instance, if organisms tend to show site fidelity in breeding, staying the next year if this year was good, and local environments are positively autocorrelated in fitness over time, this tendency helps populations persist in spatiotemporally varying environments (Schmidt, 2004). By contrast, we suspect that some organisms with environmentally forced dispersal, such as happens with wind-dispersed plant propagules and aquatic and marine organisms dispersing with water currents, may experience limited (or no) benefits of the inflationary effect.

The relationship between inflationary effects on abundance and growth is as yet undetermined. Inflated abundance in many cases implies inflated population growth at the metapopulation scale, as is illustrated by the case of persistence in a landscape of stochastic sinks. However, it is unclear whether inflated growth when rare always leads to inflated abundance. We suspect that the answer lies in how density-dependence acts on the landscape to influence how variation in fitness factors translates into long-term average abundance. Future modeling work is necessary to understand the relationships between abundance and growth rate inflation.

At this time, although the theory has been demonstrated in several laboratory experiments with model species, we do not know how widespread the inflationary effect is in nature. This is an open question for future empirical research. But the inflationary effect seems most likely to occur for species living in environments with the characteristics outlined in Table 1. The characteristics in Table 1 should be useful indicators for empirical study of the inflationary effect in nature. The inflationary effect remains a niche concept in population and community ecology, despite its apparent connection to the basic concepts of stability in metapopulation and metacommunity models. We hope that this review, and the case studies from epidemiology provided below, spark some interest in the concept, especially among empiricists.

**Table 1.** Features indicative of the inflationary effect in natural systems.

| Empirical Pattern | Link to Inflationary Effect Measure (eqn 4) |
| --- | --- |
| Nonrandom distributions of individuals across space | Spatial variation in relative density, $v_x$ |
| Temporal changes in the non-random distribution of individuals over space | Variation in local density across space and time, $v_x(t)$ |
| Spatiotemporal variation in a fitness-affecting factor | Variation in local growth rates across space and time, $r_x(t)$ |
| Undirected passive dispersal or active environmental tracking | Possibility that population is concentrated in locations and times of higher than average growth, (positive $\text{cov}(r_x(t), v_x(t))$, on average) |

### Spatiotemporal heterogeneity and infectious disease

Our emphasis thus far has been conceptual with examples from free-living protozoans and dune plants. Some others we have noted suggest application of the inflationary effect to host-parasitoid dynamics (Duncan et al., 2013), hinting at application of the inflationary effect, and the role of asynchrony and spatiotemporal heterogeneity, beyond the usual domain of the ecological sciences to epidemiology. We here argue that the inflationary effect looms large in the spread of infectious disease.

When considering infectious diseases, there are two scales at which one can envisage metapopulation dynamics. First is the metapopulation perspective of hosts as patches. From the perspective of an individual pathogen (e.g., a bacterium or virion), each individual host is an enormous habitat patch or even heterogeneous landscape (Holt, 2000). An individual SARS-CoV2 virion is about 0.1 microns in diameter and resides in a human host averaging 1.7m in height. Were we to scale up the virion to the size of its host, and re-scale host size accordingly, the host would be 29,000 km tall – more than twice the diameter of the Earth. At peak infection, the estimated viral load in a single infected human host is 10^9^ to 10^11^ viruses (Sender et al., 2021) – there are about 10^11^ stars in our galaxy.

Transmission to uninfected hosts is akin to colonization of empty patches, and pathogen clearance (or host death) is a form of local extinction for the pathogen. Susceptible-infectious-recovered (SIR) models assume hosts are either healthy (but susceptible to infection), infectious (and potentially unhealthy), or recovered from prior infection (and unable to be re-infected). This is the epidemiological analogue of the metapopulation assumption that one only need track pathogen occupancy (presence-absence) in a host habitat patch, rather than the details of within-patch (or within-host) population dynamics. To see the role of spatiotemporal heterogeneity, recognize that the usual form of SIR style models (and the constellation of similar models) assumes an exponentially distributed timing for both transmission and infection clearance at the individual level, meaning there are differences among hosts in the clearance (extinction) and infection (colonization) times. In other words, epidemiological events across hosts are asynchronous. Linking such within-host dynamics to among-host dynamics is a large challenge, with many avenues yet to be explored (e.g., analogues of emigration, see Barfield et al., 2015). This is an important theme, but here we instead go up in spatial scale, considering metapopulations of hosts (see e.g., Cliff et al. 2000).

At the level of the hosts, there can be metapopulations of hosts coupled by host movement (or possibly vector movement), which is our focus. Epidemiologists have long been concerned with aspects of disease spread in spatially distributed host populations (Ch. 7 of Keeling and Rohhani, 2007 and references therein), and with temporal variation in pathogen transmission (e.g., seasonal outbreaks in a locality; Shone et al. 2006; Moore et al. 2014). However, the impact of spatiotemporal variation along the line of our inquiry has been largely neglected.

Asynchrony among host populations in pathogen transmission can arise for many reasons. For instance, if a pathogen is transmitted by an arthropod vector along climatic gradients, seasonal peaks in pathogen transmission might develop at different times across space (Moore et al., 2014; Shone et al., 2006). This likely will be exacerbated for arthropod borne, transovarially transmitted pathogens where infected adult females may infect most, if not all, their progeny following dispersal. Spatiotemporal variation in the abundance of non-human hosts (Childs, 2004) for multi-host pathogens could influence patterns of variation in human disease.

Human behavior is often key to infectious disease transmission, and spatiotemporal variation in behavior can be a particularly potent source of spatiotemporal heterogeneous patterns of disease transmission. Public health policies also lead to spatiotemporal heterogeneity in transmission and asynchrony in control among polities or nation-states because differing levels of intensity in control measures and public health infrastructure imply infectious diseases are able to spread much more efficiently in some regions than in others. Asynchrony arises because different localities experience different patterns of human behavior over time, driven both by ‘bottom-up’ individual effects (e.g., differences among locations in social expectations about the importance of non-pharmaceutical control measures such as mask-wearing, willingness to avoid crowded venues and the like), and ‘top-down’ forces, such as imposed public health measures.

In 2020, many of the present authors noticed the potential for broad scale asynchrony in disease transmission to influence the spread of SARS-CoV-2, the etiological agent of COVID-19, early during the COVID-19 pandemic. Given that SARS-CoV-2 was a novel and virulent infectious agent, few therapeutic treatments were initially available. In the absence of known and effective medical treatment, the main approach to slow and limit the spread of SARS-Cov-2— thereby reducing stress on health care infrastructure—was the use of non-pharmaceutical interventions (e.g., masking, social distancing, closing of non-essential business places, lockdowns; hereafter NPIs). The timing, nature, and efficacy of NPIs varied across time and space as social, economic, and epidemiological conditions changed. One simple example is that government-ordered lockdowns of local non-essential businesses might respond to rises in local reported case numbers. Such feedback interacts with existing heterogeneity in the social, behavioral, economic, and demographic structure of populations to generate differential timing of lockdowns. Uncoordinated decision making about the timing of NPIs could generate substantial spatiotemporal heterogeneity in a fitness factor of SARS-Cov-2, the generalized transmission rate.

To illustrate this idea, consider a version of the model used by Kortessis et al. (2020) to explore potential consequences of asynchrony in the timing of lockdowns (see supplementary material S5 for model details). In that model, there are two patches of human hosts linked by movement. Disease spread in each patch follows the assumptions of an SIR model. All epidemiological parameters are identical between patches except for *β*(*x,t*), the generalized transmission rate, which is indexed by patch location (*x* = 1,2) and time (*t* ∈ [0,∞)). For simplicity, we assume that *β*(*x,t*) takes two values over time, *β*_0_, the baseline transmission rate, and *β*_0_(1 – *ɛ*), where *ɛ* is the effectiveness of NPIs on reducing disease spread locally in proportional terms. Furthermore, we consider that when a lockdown occurs, it persists for duration *T*, before returning to “business as usual”, with periodic bouts of lockdowns (i.e., “square-wave” temporal variation). To model differential timing of NPIs, we assume a delay between the initiation of NPIs in the different patches and keep track over how much time the two locations are in the same state (both “business as usual” or instead both under NPIs). We call the fraction of time that the two patches are in the same state Ω, the NPI overlap.

The overlap in NPI controls the magnitude of spatiotemporal heterogeneity. Indeed, the measure of spatiotemporal variance is *σ*^2^*_ST_* = (1–Ω)(*β*_0_*ɛ*/2)^2^, and on a scale proportional to the total variation is *σ*^2^*_ST_*/*σ*^2^ = (1–Ω) (see supplementary material S5). Hence, synchronized policies (Ω, = 1) do not generate spatiotemporal variability, whereas asynchronous policies (Ω < 1) do generate spatiotemporal heterogeneity. When policies are exactly out of sync (i.e., Ω = 0) spatiotemporal heterogeneity accounts for all variability in pathogen transmission.

Pairing host movement between patches with less than complete overlap in NPIs greatly elevates the rate of increase in new infections at the metapopulation scale. This is an instantiation of the inflationary effect for infectious diseases, where aspects of human behavior plausibly create spatiotemporal heterogeneity in conditions affecting the fitness of a respiratory virus. The effect is larger for more effective NPIs (i.e., larger *ɛ*, generating greater variation in transmission over time) and when the duration of application of NPIs is longer (larger *T*, details in supplementary material S5). When NPIs do not completely overlap and some infectious individuals travel between patches, a stochastic version of the model shows increasingly larger waves of infection than with complete NPI overlap (Figure 5b).

**Figure 5.**
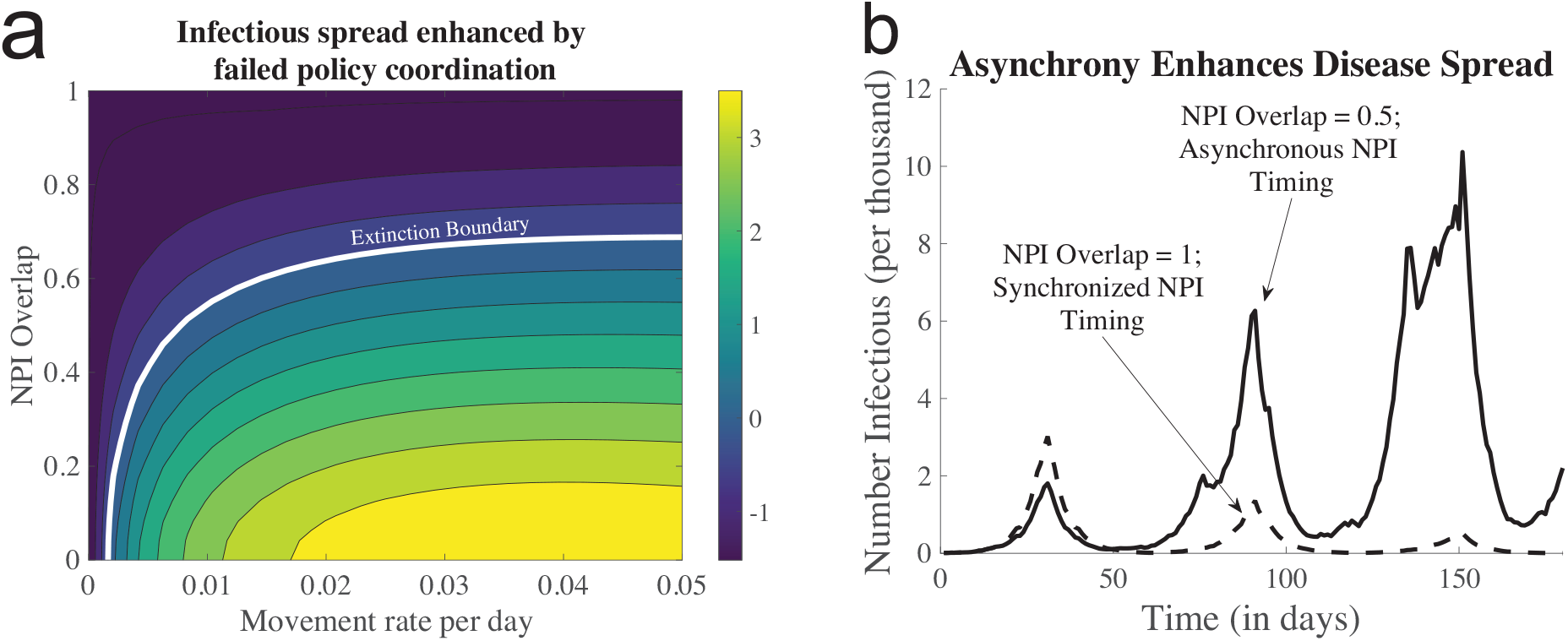
The inflationary effect in a two-patch SIR model. (a) The metapopulation-scale growth rate and how it depends on the overlap in the application of NPIs and host dispersal. (b) Example dynamics from a stochastic version of the model. Parameters: infectious duration = 4.5 days, *β*_0_ = 0.375 d^-1^ (hence *R*_0_ ≈ 1.7), *T* = 30, and *ɛ* = 0.95. In (b), *m* = 0.025 d^-1^. Further details in supplementary material S5.

Evidence from cell phone data shows human mobility changes during state-enforced NPIs in the European Union (EU) were sufficient to promote the inflationary effect in the presence of repeated, statewide lockdowns. Ruktanonchai et al. (2020) illustrated that if different countries in the EU came out of lockdown at different times (thereby generating spatiotemporal heterogeneity – as in fact was the case), then there would be many more infections and hospitalizations from COVID-19 than if states came out of lockdown in a coordinated manner. Others have modeled the timing of application of NPIs in light of this phenomenon and found it to be quite general (Berestycki et al., 2023; Eksin et al., 2021; Wang and Wu, 2022; Zhang et al., 2021). A general conclusion is that optimal local public health strategies locally depend on the public health decisions made by nearby localities. In other words, spatiotemporal heterogeneity in disease transmission, coupled with movement among locations, degraded the control of the pandemic, with serious and devastating consequences for our species across the globe.

## Discussion

Recent work on the inflationary effect clarifies and extends the intuition of empirical ecologists many decades ago about the role of spatiotemporal variation in fitness, along with dispersal, in promoting regional persistence of species. Andrewartha and Birch (1954, p. 657), for instance, remark: “A natural population occupying any considerable area will be made up of a number of … local populations or colonies. In different localities the trends may be going in different directions at the same time.” Later, they wrote admiringly of den Boer’s idea of spreading of risk (den Boer, 1968) and noted “The concept of spreading risk recognized the importance of dispersal with the natural population that enables individuals to colonize new localities” (Andrewartha and Birch, 1984, p. 175). They extensively quote den Boer (1981, p. 39), who suggests “The survival time of composite populations uninterruptedly inhabiting large and heterogeneous areas, highly depends on the extent to which the numbers fluctuate unequally in the different subpopulations.” In other words, population persistence at broad scales may rest on the interplay of dispersal and spatiotemporal variation in fitness.

The influence of spatiotemporal variation can be discerned even if populations are not found in crisply defined habitat patches. den Boer (1981) studied ground beetle ecology over 35 years and showed that though dispersal occurred, it was mainly at a low level and over short distances. The beetle *Pterostichus versicolor* fluctuated asynchronous across ten locations and did not experience extinction. By contrast, the beetle *Calathus melanocephalus* fluctuated synchronously, and did show local extinction. As Gotelli (2008, p. 96) notes for the persistent species, “… at any point in time, some populations were increasing in size and acting as source populations that prevented the extinction of other, declining sink populations.” The other species experienced synchrony in its fluctuations, so there were times when conditions were uniformly bad for it everywhere. The data den Boer (1981) presents resembles qualitatively the patterns shown in the simple metapopulation model of Figure 1.

A new generation of empirical research is needed to establish how widespread the inflationary effect is in nature (see Table 1 for the necessary features), both in single species metapopulation dynamics and in metacommunities where species interactions may contribute to fluctuations in fitness over space and time.

### Invasion biology

While initial studies of the inflationary effect highlighted persistence of a metapopulation of sinks (Roy et al. 2005), a novel phenomenon highlighted in Kortessis et al. (2020) is that the inflationary effect could also speed up the course of an invasion, even if metapopulation persistence is not an issue. We suggest that this effect might contribute to a well-known phenomenon in invasion biology – biphasic and accelerating patterns of expansion for invasive species, including initial lag phases (Hui and Richardson, 2017, pp. 28–37 for many examples). In the case of the common starling, *Sturnus vulgaris*, for instance, its spread was sluggish in the first few decades after its introduction into North America (Okubo, 1988), and then it sped up. Similarly, West Nile Virus was slow to disperse throughout the eastern USA and remained at low incidence in the first couple years following introduction in New York City in 1999, but then rapidly expanded in 2002 (Petersen and Hayes, 2008).

A variety of explanations for this pattern have been suggested (e.g., transitions among different habitats, Okubo 1988; stratified diffusion, Shigesada et al. 1995). The inflationary effect provides another viable explanation. If there are *n* locations with independently fluctuating growth rates, and a species is introduced into one of those locations at random and thereafter disperses, then its initial expected growth rate (prior to much dispersal) is simply the arithmetic average growth rate over those locations. However, as time goes on and individuals move among locations, then the covariance term in expression (4) grows, and if this is positive, the invasion will accelerate. Spatial population data, especially for invasive insects, could be used to assess the role of the inflationary effect during invasion.

### Natural enemy-victim interactions

Effective predators tend to overeat their prey, and then suffer extinction themselves. The ‘spreading of risk’ among different populations, loosely coupled by dispersal, is a likely reason for the persistence of strong natural-enemy interactions. In the classic experiments of Huffaker (1958), the persistence of a predatory mite (*Typhlodromus occidentalis*) with a prey mite (*Eotetranychus sexmaculatus*) in a mesocosm comprised of an array of oranges depended on limited dispersal, and the fact that dynamics were asynchronous across space. This experiment inspired an experiment with a bruchid beetle *Callosobruchus chinensis*, feeding on beans, attacked by a pteromalid parasitoid *Anisopteromalus calandrae* (Hassell and May, 1988). In an environment without spatial structure, the parasitoid rapidly overexploited its host, then went extinct (allowing the host to rebound). When each beetle was put in its own patch, with restricted dispersal among patches, the host-parasitoid interaction persisted indefinitely. Here, the physical environment was homogeneous (by experimental design), and spatiotemporal variation in each species’ fitness arose entirely from its localized interaction with the other species and its own direct density dependence.

It is likely that this effect explains the robust persistence of a wide range of strong natural enemy-victim systems (Hassell, 2000). Because dispersal is often localized, the asynchrony needed for spatiotemporal variation to be expressed depends upon spatial scale. The relevant scale(s) of temporal fluctuations will vary with the generation time, demography, and movement rates of the focal species (Bernhardt et al., 2020). For example, Wilson et al. (1998) examined how the persistence of host-parasitoid and host-parasitoid-hyperparasitoid interactions depends upon the size of the spatial lattice within which these dynamics played out. They found that food chains of strongly interacting specialist species require larger arenas to persist. Adjacent areas are more likely to be synchronous than are distant, far-flung locations. Island and patch size effects on food chain length of specialists might thus reflect in part the importance of spatial scaling of asynchrony in population dynamics among localities within defined spatial arenas (Holt, 2010; e.g., islands, habitat fragments, Wilson et al., 1998).

### Interspecific competition

The inflationary effect has consequences for competitive interactions and species coexistence. A local habitat might be a sink for a species not because the abiotic environment is poor but because of the presence of a superior competitor. Inferior species can nonetheless persist because of recurrent immigration from a source. Temporal variability in local growth conditions could inflate the abundance of the inferior competitor, thus increasing its impact on the superior competitor. Long et al. (2007) showed experimentally in a laboratory microcosm using bacterivorous protists that this effect could sometimes lead to the local extirpation of the locally superior (on average) species.

The inflationary effect may also lie at the foundation of a classic mechanism of coexistence in metacommunities: the competition-colonization tradeoff. Classic models (e.g., Tilman, 1994) show that with strongly asymmetric competition, multiple species can coexist provided there are external sources of extinction that wipe out superior competitors. This requires however that patches be colonized asynchronously. If the superior species is a sluggish colonizer, there is a time period during which fitness of the subordinate in the empty patch once it arrives is high, and far distant locations are unlikely to be colonized simultaneously by the superior species. Once the superior species arrives, however, the fitness for the subordinate is low, and it stays low until the superior species suffers extinction. Thus, the dynamics of the superior species in effect generates a positively autocorrelated environment for the fitness of the inferior species. This can promote coexistence, provided there is asynchrony among patches in extinction and subsequent recolonization (Roy et al., 2005).

The inflationary effect also underpins the spatiotemporal dynamics of the spatial insurance hypothesis (Loreau et al., 2003; Shanafelt et al., 2015; Thompson et al., 2017). This hypothesis posits that biodiversity provides stability to ecosystem functioning across a metacommunity. This effect stems from the interaction between dispersal, differences among competing species in their use of the environment, and spatiotemporal environmental fluctuations. If different metacommunities experience distinct temporal patterns of environmental conditions, species differentially adapted to the environment are expected to thrive in any metacommunity (as a temporary source) for a finite period before the environmental state of the patch switches. This results in spatiotemporal shifts in the source or sink status of patches across the metacommunity. Dispersal ensures that the species adapted to the new environmental conditions locally are available to replace less adapted species as the environment changes. As a result, biodiversity may enhance and buffer ecosystem processes by virtue of periods of inflationary growth and spatial exchanges among local ecosystems, even when such effects do not occur in a closed homogeneous system. Factors that synchronize population dynamics across ecosystems (Reuman et al., in press) or constrain dispersal will erode the multispecies inflationary dynamics that contribute to metacommunity stability.

### Genetic variation and evolutionary dynamics

Our focus has been on ecological consequences of spatiotemporal variation and the inflationary effect, but there are doubtless evolutionary effects, as well. Here, we mention just a few examples. The model of Loreau et al. (2003), which focuses on species, can be interpreted as a model of asexual selection maintaining genetic diversity in a spatiotemporal environment (Levene, 1953). Wieczynski and Vasseur (2016) demonstrated that the inflationary effect could promote the maintenance of intraspecific genetic variation when different genotypes are differentially superior at distinct times, and showed that this could increase overall population size and promote population persistence. Expression (4) describing the inflationary effect in terms of the covariance between local growth rate and relative abundance can be viewed as a component of fitness that selection can act upon. If a behavioral variant arises that increased the match between the spatial distribution of that variant, and its temporarily greater local growth rate, that variant would have a selective advantage. This is one way of describing the fitness advantage of habitat selection, a point illustrated by Altenberg (2012) in the context of the evolution of dispersal.

The inflationary effect has been shown to promote the persistence and evolution of more contagious viral strains. In essence, patches with high transmission rates foster genetic diversity in the pathogen population. New variants can then spread to areas with low transmission rates and establish themselves in communities where other, less infectious strains may not persist. Overall, this creates an environment that both promotes the evolution of new variants, and the persistence of highly contagious variants in areas where community transmission could otherwise be stopped, as suggested in Europe through the early stages of the COVID-19 pandemic (Lemey et al., 2021). In the winter of 2020, the B.1.1.7 variant was suspected to first appear in the United Kingdom and Spain, where intervention measures were mostly lifted. Nonetheless, this variant was able to rapidly spread across the continent and persist in countries still undergoing lockdown measures because of its increased contagiousness.

### Conservation and management

Drivers of the inflationary effect can be leveraged to inform conservation and management decisions. The greatest threats to biodiversity --- global change, habitat destruction, overexploitation, among others (Bellard et al., 2022; Brook et al., 2008) --- could act to increase or decrease the strength of inflationary effects operating in nature. For instance, enhanced weather variability could decrease the spatial synchrony of local population growth (Hansen et al., 2020), thus increasing the inflationary effect. That said, increased incidence of extreme climatic events (e.g., droughts and heatwaves) at large spatial scales could synchronize populations (a generalized Moran effect; Hansen et al. 2020, see also Reuman et al. in press), thereby weakening the inflationary effect.

Scientists already recognize the importance of incorporating metapopulation processes for pursuing conservation priorities (Akçakaya et al., 2007), but considering the inflationary effect emphasizes the critical stabilizing nature of metapopulations. Facilitating movement between local populations (e.g., facilitated by corridors) is well established as beneficial for conserving species at risk of extinction. However, the inflationary effect suggests that too much movement might degrade asynchronous population fluctuations (to at least some degree). Furthermore, the timing of reintroduction or restocking of populations, might be more effective if it maintains or generates asynchronous dynamics.

Similar ideas can be applied to management of fire-obligate communities, the complexity of which requires a carefully planned decision-making framework (Kelly et al., 2015). Managed burning regimes might be made more effective by thinking about the heterogeneity or homogeneity in burns created across the landscape by land managers, which may affect decisions about the acreage per burning event, the timing and spatial arrangement of burns, and burn intensity. All such factors influence the spatiotemporal heterogeneity affecting the dynamics of species in fire-prone systems that might benefit (or rely) on the inflationary effect. Similar concepts apply in fisheries. Prescriptions about quotas might include considerations for how they promote or degrade asynchrony. Lack of these considerations might foster intense fishing on a few populations with high numbers, which synchronizes population dynamics across large scales, reducing the so-called population portfolio (Stier et al., 2020) and eroding stability provided by the inflationary effect.

### Infectious disease and epidemiology

While we have emphasized the inflationary effect for the spread of SARS-CoV-2, it likely plays a role for many other pathogens, such as measles. Prior to vaccination in the late 1960s, measles had biennial epidemics that were synchronized throughout England and Wales. Bolker and Grenfell (1996) documented how widespread vaccination eliminated large epidemics, which were the putative causative synchronizing factor, and so decoupled measles dynamics across space. Such asynchrony was hypothesized to reduce the chances of extinction of measles across all geographies simultaneously. Indeed, despite consistent declines in incidence of measles for three decades after vaccine introduction, measles incidence is far above elimination and eradication targets (Winter et al., 2022).

Bubonic plague (caused by the bacterium *Yersinia pestis*), which occasionally causes localized epidemics in humans, appears to be maintained by the inflationary effect in rat populations (Keeling and Gilligan, 2000). Locally abundant susceptible populations of rats (wherein the pathogenic bacterium’s fitness is high) temporarily cause large outbreaks that spill over and infect humans. Epidemics in the rat population happen differently across space and are localized in time, leading to a patchwork of populations in the landscape with differential fitness from the perspective of the pathogen --- with knock-on consequences for human health.

### A final thought

This paper is an invited contribution to a special feature of The American Naturalist on the theme of neglected and misunderstood ideas in ecology. One of the lessons to emerge from the recent pandemic is that governmental policymakers must seriously confront the complicated dynamical issues we bring out here. However, the situation is not unique to the recent pandemic. Pathogen eradication is extremely rare among human pathogens, a fact we believe occurs in part because of the large heterogeneities present at the scale of human societies. For emerging infectious diseases, the lesson is to address exponentially growing processes quickly and aggressively before they get out of hand. Beyond this, we submit that ignorance of the emergent effects of spatiotemporal variability in local growth rates, given dispersal among locations, contributed substantially to the failure of governmental control of the Covid pandemic in its early phases in many parts of the globe. In New York, for instance, there was a ‘block-by-block’ control strategy (Guarino, 2020). In the United Kingdom, then Prime Minister Boris Johnson emphasized local lockdowns and local ‘whack-a-mole’ strategies (BBC 2020).

This absence of coordinated control efforts, at spatial scales from local municipalities up to entire continents, given that there was continued movement of asymptomatic, infectious individuals among locations—estimated to be responsible for up to 35% of all infections (Sah et al., 2021)—likely contributed to the pervasive failure of control. Failure to recognize the population dynamical consequences of spatiotemporal variability present in landscapes, with movement across those landscapes, perhaps fostered a great deal of human suffering. One can only hope that nations, and indeed the world, learned something from this experience.

## Acknowledgements

We thank Nick Gotelli, Rachel Germain, and Sebastian Schreiber for useful discussions. NK, MWS and RDH thank NSF grants for support, and RDH thanks the University of Florida Foundation for continued support. AG is supported by the Liber Ero Chair in Biodiversity Conservation and an NSERC Discovery Grant.

## Data Availability Statement

Code to reproduce the figures here and in the supplementary material can be found in the Github repository: https://github.com/kortessis/Asynchrony-and-Inflationary-Effect. The code will be archived in Zenodo pending acceptance of the manuscript for publication, following any requested changes by reviewers and editors.

## Footnotes

1 While this is often overlooked in more contemporary descriptions, environmental heterogeneity was at the forefront of Levins’ mind when developing his model. Levins’ original article articulating the metapopulation model is titled “Some demographic and genetic consequences of environmental heterogeneity for biological control”, the wording of which highlights the influence of environmental heterogeneity in Levins’ thinking.

