## Supplementary Information for "Metapopulations, the inflationary effect, and consequences for public health"

### S1 Formalizing Spatio-Temporal Variation

Here we provide a mathematical justification for the expressions of different variance components.

We begin with a random variable $F(t,x)$ that can be indexed by time $t$ and spatial location $x$. To calculate its variance, and moreover to break that variance into different components, we need to introduce the conditional variables

$$F|x \tag{S1}$$

and

$$F|t. \tag{S2}$$

The variable $F|x$ gives possible range of outcomes across time at a specified location $x$. In essence, it is the distribution of conditions possible in time in location $x$. Similarly, $F|t$ gives the possible range of outcomes (and their associated probabilities) across space at particular time $t$.

We can calculate the variance in $F(t,x)$ using the law of total variance by conditioning on spatial location such that

$$\mathrm{Var}(F) = E_x[\mathrm{Var}_t(F|x)] + \mathrm{Var}_x(E_t[F|x]), \tag{S3}$$

where $F|x$ is the random variable $F$ in a particular location $x$, $E_x[\cdot]$ and $E_t[\cdot]$ are spatial and temporal expectations, respectively, and $\mathrm{Var}_x(\cdot)$ and $\mathrm{Var}_t(\cdot)$ are spatial and temporal variances, respectively.

To partition this variation into space, time, and spatio-temporal components, we introduce the concepts of the temporal mean field and the spatial mean field. A temporal mean field model is one in which conditions that vary over time are represented by their mean. A model of the temporal mean field includes spatial variation but any temporal variation is represented simply by the mean. Let $\tilde{F}_x := E_t[F|x]$ be the distribution of fitness-affecting conditions present in such a model. These conditions have variance

$$\sigma_S^2 = \mathrm{Var}_x(\tilde{F}_x) = \mathrm{Var}_x(E_t[F|x]), \tag{S4}$$

which is entirely comprised of spatial variation and so represents "spatial-only" variation.

A spatial mean field model is one in which conditions that vary across space are represented by their mean. A model of the spatial mean field includes temporal variation but any spatial variation is represented simply by the mean. Let $\bar{F}_t := E_x[F|t]$ be the distribution of fitness-affecting conditions present in such a model. These conditions have variance

$$\sigma_T^2 = \mathrm{Var}_t(\bar{F}_t) = \mathrm{Var}_t(E_x[F|t]), \tag{S5}$$

which is entirely comprised of temporal variation and so represents "temporal-only" variation.

Substituting (S4) into (S3) and adding and subtracting $\sigma_T^2$ yields

$$\mathrm{Var}(F) = E_x[\mathrm{Var}_t(F|x)] + \mathrm{Var}_x(E_t[F|x]) \tag{S6}$$

$$= E_x[\mathrm{Var}_t(F|x)] - \mathrm{Var}_t(E_x[F|t]) + \sigma_S^2 + \sigma_T^2 \tag{S7}$$

$$\mathrm{Var}(F) = \sigma_S^2 + \sigma_T^2 + \sigma_{ST}^2, \tag{S8}$$

where in the last line we have defined $\sigma_{ST}^2 = E_x[\mathrm{Var}_t(F|x)] - \mathrm{Var}_t(E_x[F|t])$ as the magnitude of spatio-temporal variation, and is that variation in $F$ which remains after accounting for spatial-only and temporal-only variation.

### S2 Stochastic Metapopulation Model

Here we provide a mathematical overview of the metapopulation model with (patch-level) demographic stochasticity. This model is conceptually similar to the traditional metapopulation model as introduced by Levins, but follows a finite number of patches. Given the finite number of patches, we must deal with the metapopulation analogue to demographic stochasticity, which is random deviations in extinction and colonization from large-scale averages.

The model is a discrete-time Markov chain for $n$ patches, and it models the occupancy of a all patches as denoted with the time-dependent vector $\mathbf{O}(t) = (O_1(t), O_2(t), \ldots, O_n(t))^T$. Each $O_i(t)$ can be either 0, corresponding to an empty patch, or 1, corresponding to an occupied patch. Occupancy changes states (or stays in the same state) with extinction and colonization probabilities, $E_i$ and $C_i(t)$, respectively. To formalize the model, a typical approach is to write that the transition probabilities for all patches. To do so, define $P_t^i$ as

$$P_t^i(a,b) := P\Big(O_i(t+\Delta t) = b | O_i(t) = a\Big), \tag{S9}$$

the probability that patch $i$ in state $a$ at time $t$ transitions to state $b$ in the time interval $(t, t+\Delta t]$. Given the extinction and colonization probabilities above, we have

$$P_t^i(0,0) = 1 - C_i(t) \qquad P_t^i(0,1) = C_i(t) \tag{S10}$$

$$P_t^i(1,0) = E_i(t) \qquad P_t^i(1,1) = 1 - E_i(t). \tag{S11}$$

Given that we are only interested in illuminating the consequences of asynchrony, we keep the assumptions of the model simple. In particular, we assume that all patches are the same size and are equally connected, which corresponds to the assumptions developed initially by Levins for the first metapopulation model in continuous-time. We set

$$E_i = 1 - \exp(-e\Delta t) \tag{S12}$$

for all patches, where $e$ is the infinitesimal rate of extinction, and $\Delta t$ is a small unit of time. This model of extinction has the typical biological interpretation that all patches are effectively identical with respect to factors that influence extinction, and corresponds to an extinction rate of $e$ in the limit as $\Delta t \to 0$.

We also assume that the probability that an unoccupied patch stays unoccupied is

$$1 - C_i(t) = \exp\big( - cP(t)\Delta t \big), \tag{S13}$$

such that the colonization probability is

$$C_i(t) = 1 - \exp\big( - cP(t)\Delta t \big). \tag{S14}$$

This colonization model implies that colonization rates depends on the occupancy on the landscape and occur at infinitesimal rate $cP(t)$. Indeed, in the limit as the time interval becomes very small, this discrete-time model converges on the continuous-time metapopulation model (1) of the main text that was originally presented by Levins. In our simulations, we chose $\Delta t$ = 0.0005.

The conditional transition probabilities (S10)-(S11) alone are not enough to specify this model. For finite patch numbers, we must also specify the probability distribution for number of extinction (and colonization) events each change in time, $\Delta t$. For example, with 5 occupied patches and a 50% probability that each patch goes extinct in one unit of time, there is a small, but finite chance that none go extinct in the time period of $\Delta t$ and a similar small but finite chance that all go extinct over the same time range.

A typical probability distribution chosen for this is the binomial distribution such that the number of extinctions is $\text{Binom}(\sum_i O_i(t), E_i)$. However, this distribution makes the strong assumption that the extinction event for one patch is independent of that for any other patch. Biologically, this implies that the factors influencing extinction in one patch at one time are independent to those in any other patch across space. That may very well be true, but is a specific case of a broader set of possibilities.

To model the broader range of possibilities, we use a latent variable approach that describes a hypothetical environmental variable affecting extinction and colonization in each patch. We then use properties of the extinction and colonization probabilities to create a reaction norm relating local environmental characteristics to extinction or colonization events.

Note that while this works for discrete-time models, correlated events can be described in true continuous-time as correlated times to extinction. However, such a mathematical description is beyond the scope of this paper. The approach here is sufficient to illustrate the point.

To model correlated events, let $X_i(t)$ be the environmental conditions in patch $i$ at time $t$. For simplicity for modeling correlations between variables, we assume that $X_i(t)$ is a standard normal random variable that is i.i.d. (independent and identically distributed) over time; hence, $X_i(t) \overset{\text{iid}}{\sim} \mathcal{N}(0,1)$ for all $i$. To include correlations between conditions in patches across space, we collect all individual environmental variables as a multivariate collection $\mathbf{X}(t) = (X_1(t), X_2(t), \ldots, X_n(t))^T$ that follows a multivariate normal with zero mean and covariance matrix $\Sigma$. Because all the marginal variables are standard normal, $\Sigma$ has values of 1 for all diagonal elements and therefore takes the interpretation of a correlation matrix. Again, for simplicity, we assume this correlation matrix has the a simple structure where $\text{Corr}(X_i(t), X_j(t)) = \rho$, for all $i \neq j$. Written succinctly,

$$\mathbf{X}(t) \overset{\text{iid}}{\sim} \mathcal{N}_n\Big(\mathbf{0}, \Sigma = \rho\mathbf{1}\cdot\mathbf{1}^T + (1-\rho)\mathbf{I}\Big), \tag{S15}$$

where $\mathbf{1}$ is an $n$-length column vector of ones. Correlations, in general, take values between -1 and 1. However, given that the correlation applies to all possible pairs of patches, the multivariate normal will not permit $\rho$ near -1 when $n > 2$. Indeed, larger $n$ restricts the range of permissible values of $\rho$ based on the requirement that $\mathbf{\Sigma}$ be positive semi-definite. Any textbook on multivariate statistics will cover such a restriction. We only consider $\rho$ values between 0 and 1.

To translate this environment into an extinction (or colonization) event, we convert the environmental variables, $\mathbf{X}(t)$ into multivariate Bernoulli random variables using copulas. Copulas are used to create correlated random variables in cases where the marginals are of different distributional form or more generally where the multivariate distribution has no known analytical form. Copulas and their use in ecology are discussed by Ghosh et al. (2020). For examples of the use of copulas for questions of life history evolution in plant communities, see Kortessis and Chesson (2018, 2021).

Here, the goal is to have a multivariate distribution of extinction events at each time step under the provision that each marginal distribution has the probability $E_i$ of an extinction event in time $t$. Thus, we need a random variable

$$Y_i(t) = \begin{cases} 1 & \text{with probability } E_i \\ 0 & \text{with probability } 1 - E_i \end{cases}, \tag{S16}$$

where $Y_i(t) = 1$ means that a population $i$ that was occupied in time $t$ goes extinct in time $t+1$.

The transformation for $Y_i(t)$ from $X_i(t)$ is

$$Y_i(t) = \begin{cases} 1 & \text{if } \phi(X_i(t)) < E_i \\ 0 & \text{if } \phi(X_i(t)) \geq E_i \end{cases}, \tag{S17}$$

where $\phi(\cdot)$ is the standard normal cumulative distribution function. Hence, $\phi(X_i(t)) \in (0,1)$. Moreover, $\phi(X_i(t))$ has a uniform distribution (indeed, all continuous random variables are uniformly distributed when transformed with their distribution function, a fact that can be found in most graduate textbooks in probability; e.g., Theorem 2.1.10 in Casella and Berger 2001). Figure S1 illustrates how a random sample from a normal distribution (red points), once transformed through its distribution function (solid line), yields a random sample from a uniform distribution (purple points).

Because $\phi(X_i(t)) \sim \text{Uniform}(0,1)$, $P(\phi(X_i(t)) < E_i) = E_i$, and $P(\phi(X_i(t)) \geq E_i) = 1 - E_i$. Hence, $P(Y_i(t) = 1) = E_i$ and $P(Y_i(t) = 0) = 1 - E_i$, meaning $Y_i(t) \sim \text{Bernoulli}(E_i)$. In our

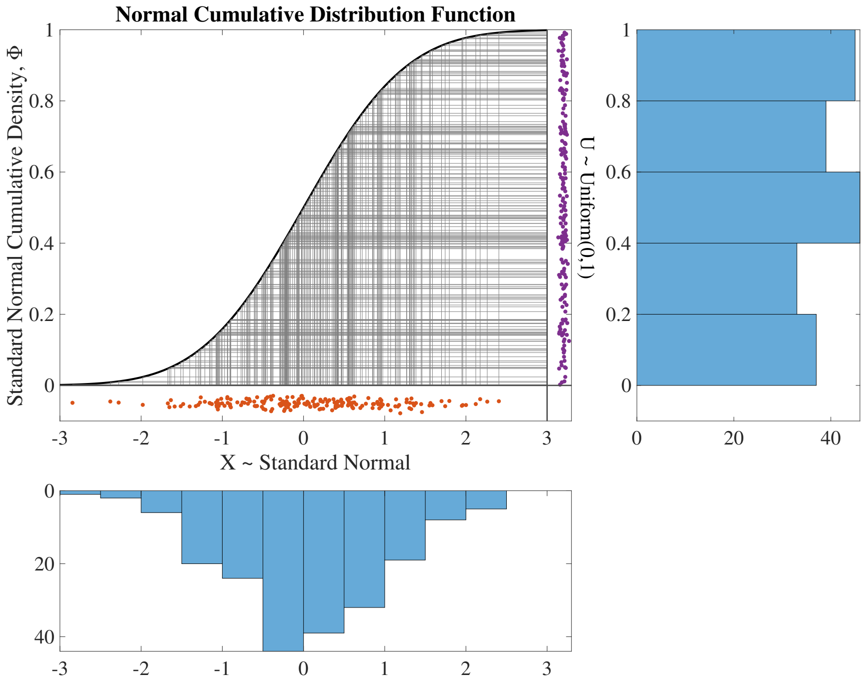


Figure S1: Demonstration of the fact that a sample from a standard normal distribution (red points in main figure; frequency distribution shown in the lower panel) can be transformed to a random sample from a uniform distribution (purple points; frequency distribution shown in the right panel). The transformation is done by the normal cumulative distribution function, which is the curve in the main panel. Gray lines show the mapping of a random sample from the standard normal through the distribution function to yield a value between 0 and 1.

simulations, we sample $\mathbf{X}(t)$ to calculate $\mathbf{Y}(t)$ according to equation (S17), and then use $\mathbf{Y}(t)$ to determine which local populations go from occupied to unoccupied from time step $t$ to time step $t + \Delta t$.

Importantly, the correlation structure embedded in the multivariate distribution $\mathbf{X}(t)$ is likewise embedded in the multivariate distribution of extinction events, $\mathbf{Y}(t) = (Y_1(t), Y_2(t), \ldots, Y_n(t))$, because $Y_i(t)$ is a monotonic function of $X_i(t)$.

Figure S2 shows a version of this model for 2 patches where the correlation is very strong in the environmental variables (Fig. S2a) such that, when the latent variables are transformed to extinction events (Fig. S2b), the spatio-temporal pattern of extinction events shows strong synchrony over space (Fig. S2c).

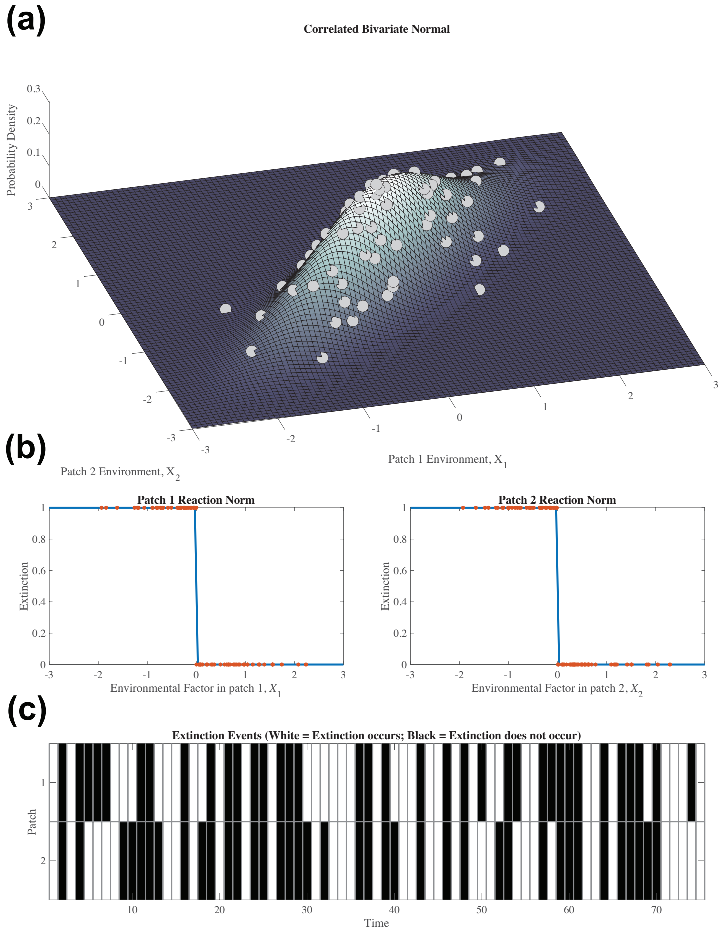


Figure S2: Example construction of correlated extinction events with two patches in the occupancy model. (a) The correlated probability distribution for $\mathbf{X}$ representing the two environmental factors present in the patches that affect extinction. Each marginal has mean 0 and unit variance, and the correlation between marginals is $\rho_E = 0.85$. The marginal variables can be considered as latent variables measuring habitat quality. Gray circles represent a random sample from this bivariate distribution. (b) The environmental factors are translated to extinction events in each patch. The relationship between $X_i(t)$ and the extinction event is given by (S17) and is shown by the blue line with $e = 0.5$. The random sample in (a) is shown by the red points in (b). (c) The spatio-temporal pattern of extinction events from the random sample in (a) and transformed in (b). Because of the correlated nature of the environmental variables, extinction events are also correlated across space such that an extinction event in one patch means that extinction is likely in the other patch.

##### *S2.1 Simulation Details*

To simulate extinction times and mean occupancy with this model, we chose a population with $n = 50$ patches. Each occupied patch has extinction probability, $E_i(t) = 1 - \exp(-e\Delta t)$, with $e = 0.25$. Since we set $\Delta t = 0.005$, the expected time to extinction is $\Delta t / E_i(t) \approx 4$ units of time. We also chose $c = 0.5$ such that colonization probability of any unoccupied patch in a small time step is $C_i(t) = 1 - \exp(-cP(t)\Delta t)$. Under this model, when $P(t) = 0.5$, patches have an expected time unoccupied also of $\approx 4$ units of time. At this point, extinction and colonizations equal each other in the large patch number limit, meaning $P(t) = 0.5$ is the equilibrium occupancy for the deterministic form of the model.

For each simulation, we begin with 13 of the 25 patches initially occupied, i.e., $O_i(t = 0) = 1$ for $i = 1, 2, \ldots, 13$ and $O_j(t = 0) = 0$ for all $i \neq j$. Each time step, we randomly sample $\mathbf{X}(t)$ and calculate $\mathbf{Y}(t)$ for each patch. If a patch is occupied and $Y_i(t) = 1$, then the patch becomes unoccupied in the next time step. For unoccupied patches in time $t$, we take a random sample from a Binomial random variable with $n(1 - P(t))$ trials (i.e., the number of unoccupied patches) and "success" probability $C(t)$. Sampling from a Binomial distribution implies independence of colonization events across unoccupied patches, a simplifying assumption made here for simplicity. Non-independence of extinction is sufficient to demonstrate the effect of spatio-temporal heterogeneity.

We repeated this process for 100 units of time (i.e., $100/\Delta t = 20{,}000$ time steps) and evaluated whether the metapopulation had gone extinct (i.e., $P(t) = 0$). If it had, we recorded the first time of metapopulation extinction. If it had not yet gone extinct, we continued for another 100 units of time, repeating this process until the metapopulation had gone extinct.

We repeated this process for 100 replicate sample paths of extinction and colonization events for a single value of $\rho_E$. Extinction times are approximately log-normally distributed across the replicate simulations, so we plotted extinction times on the $\log_{10}$ scale.

#### S3 Inflated Abundance in a Source-Sink Model with Spatio-Temporal Heterogeneity

Here we provide analytical justification for the inflationary effect in a model with a fluctuating sink with constant immigration.

Let $N(t)$ be the density of individuals in the sink at time $t$ ($t = 0, 1, 2, \ldots$). We assume that the population grows exponentially in the sink with long-term growth rate $\bar{r} < 0$, such that the patch is better described as a *stochastic sink*. But the growth rate in any given of time $(t, t+1]$ is given by $r(t)$. Finally, we assume that there is a constant source of immigrants that enter the sink. The number that enter in a given time interval per area of the sink is $I$.

The population size in time $t+1$ is then given by the following equation:

$$N(t+1) = N(t)e^{r(t)} + I. \tag{S18}$$

The per-capita growth rate for this model is

$$g(r,N) \equiv \ln N(t+1) - \ln N(t) = \ln \{e^{r(t)} + I/N(t)\}. \tag{S19}$$

This equation allows to make some observations about the qualitative nature of the dynamics in the model. The first is that the population is bounded from above because

$$\lim_{N\to\infty} g(r,N) \to r(t) \tag{S20}$$

which, on average, is negative because $r(t)$ is negative on average. Thus, the population has a tendency to decline when its density is too high.

Moreover, we know that population size is bounded from below because

$$\lim_{N\to 0} g(r,N) \to \infty, \tag{S21}$$

and as such, the population has a strong tendency to recover as it becomes rare. This is the strong stabilizing effect of immigration.

Taken together, these two observations suggest that the population stays within some range of population sizes and does not increase to infinity, nor decline to zero. Hence, at some temporal scale, the average per-capita growth rate is 0, i.e., $E[g(r,N)] = 0$.

This equilibrium is straightforward to define in the case of a constant environment, i.e., $r(t) = \bar{r}$ for all $t$. Equilibrium population size, $N^*$, satisfies the equation

$$g(r,N^*) = 0 \Rightarrow e^r + I/N^* = 1. \tag{S22}$$

The solution to (S22) is $N^* = I/(1-e^r)$, which is the equilibrium population size in a constant environment.

By nature of the fact the population is bounded from below and bounded from above by the arguments above (provided $E[r(t)] < 0$), then it follows that, in the long-term, the population neither grows nor declines, on average. Hence, the long-term growth rate of the population is $E[g(r(t),N(t))] = 0$. A small-variance approximation of $g$ around the constant environment case (i.e., $r^* = \bar{r}$ and $N^* = I/(1-e^{\bar{r}})$), yields the following equation for the growth rate at time $t$:

$$\begin{aligned} g(r,N) = g(\bar{r},N^*) + e^{\bar{r}}(r-\bar{r}) - \frac{(1-e^{\bar{r}})^2}{I}(N-N^*) + \frac{1}{2}\frac{e^{\bar{r}}}{1-e^{\bar{r}}}(r-\bar{r})^2 \\ + \frac{1}{2}\frac{(1+e^{\bar{r}})(1-e^{\bar{r}})^3}{I^2}(N-N^*)^2 + o(\sigma^2), \end{aligned} \tag{S23}$$

where we assume that $r-\bar{r} = O(\sigma)$ and $N-N^* = O(\sigma)$, $\sigma$ small.

Noting that $g(\bar{r}, N^*) = 0$, and taking the expectations of the left and right hand sides of (S23) yields

$$E[g(r,N)] = -\frac{(1-e^{\bar{r}})^2}{I} E[(N - N^*)] + \frac{1}{2}\frac{e^{\bar{r}}}{1-e^{\bar{r}}}\text{Var}(r) + \frac{1}{2}\frac{(1+e^{\bar{r}})(1-e^{\bar{r}})^3}{I^2}\text{Var}(N) + o(\sigma^2). \quad \text{(S24)}$$

Since $E[g(r,N)] = 0$, simple rearrangement of (S24) yield the following approximate difference between the average population size and the equilibrium population size under constant conditions:

$$E[N] - N^* \approx \frac{1}{2}\left[\frac{Ie^{\bar{r}}}{(1-e^{\bar{r}})^3}\text{Var}(r) + \frac{(1+e^{\bar{r}})(1-e^{\bar{r}})}{I}\text{Var}(N)\right] \geq 0, \quad \text{(S25)}$$

where the approximation sign denotes $o(\sigma^2)$, and the inequality follows from the fact that all the terms on the right hand side of the approximation are non-negative provided $\bar{r} < 0$. Hence, $E[N] \geq N^*$, meaning that the average population size under variable local growth rates is larger than the model under constant growth rates. This is simple example of the inflationary effect for small variation in local growth rates.

#### *S3.1 Quantifying the magnitude of spatio-temporal heterogeneity*

The source-sink model has spatio-temporal heterogeneity because the sink's growth rates fluctuate while it is implied that the source's growth rates do not. Constant immigration to the source is most consistent with a constant fraction of the population in the source that leaves each time step. This also requires that the population size in the source be at an equilibrium

By making the dynamics of the source patch explicit, we can quantify the spatio-temporal variability $\sigma^2_{ST}$.

Label the source as patch 1 and the sink as patch 2. Assume that the source has density-independent growth rate $r_1$, a constant, that is reduced by density-dependence with per-capita strength $\alpha$. Following growth, a fraction $d$ of individuals disperse to the sink. With these assumptions, the dynamics in the sink follow

$$N_1(t+1) = N_1(t)e^{r_1 - \alpha N_1(t)}(1-d), \quad \text{(S26)}$$

and the total number of dispersers to the sink in one unit of time is $I(t) = N_1(t)\exp(r_1 - \alpha N_1(t))d$. Immigration becomes constant when the source population reaches equilibrium, $N_1^*$, can be found by solving

$$N_1^* = N_1^* e^{r_1 - \alpha N_1^*}(1-d), \quad \text{(S27)}$$

and has the non-trivial solution

$$N_1^* = \frac{r_1 + \ln\{1-d\}}{\alpha}. \quad \text{(S28)}$$

Note that because $0 \leq d \leq 1$, $\ln\{1-d\} \leq 0$, meaning that $r_1 > \ln\{1-d\}$ in order for the source population to have a positive equilibrium density.

At this equilibrium, the growth rate prior to dispersal in the source is $\exp(r_1 - \alpha * N_1^*) = 1/(1-d) \geq 0$, which is a constant. The number of immigrants to the sink at equilibrium can also be calculated as

$$I = d\exp(r - \alpha N_1^*)N_1^* = \frac{d}{1-d} \cdot \frac{r_1 + \ln(1-d)}{\alpha} \tag{S29}$$

With the growth rates in hand, we can define the fitness factor as the local growth rates themselves, which are

$$\begin{aligned} F(t,1) &= -\ln(1-d) \\ F(t,2) &= r(t). \end{aligned} \tag{S30}$$

Given that the growth rate in source is constant, the temporal mean is $E_t[F|x=1] = -\ln(1-d)$ and the temporal variance is $\text{Var}_t[F|x=1] = 0$. The temporal mean and variance for the sink growth rates are $E_t[F|x=2] = \bar{r}$ and $\text{Var}(F|x=2) = \sigma_r^2$. Since both patch types are equally common (i.e., there is one of each), we weight their contribution to growth equally in the spatial variance, which is

$$\begin{aligned} \sigma_S^2 &= \text{Var}_x(E_t[r|x]) \\ &= \frac{(-\ln(1-d) - \bar{r})^2}{4}. \end{aligned} \tag{S31}$$

The temporal variance is calculated as

$$\begin{aligned} \sigma_T^2 &= \text{Var}_t(E_x[r|t]) \\ &= \text{Var}_t\left(\frac{-\ln(1-d) + r(t)}{2}\right) \\ &= \text{Var}_t\left(\frac{r(t)}{2}\right) \\ &= \frac{\sigma_r^2}{4}. \end{aligned} \tag{S32}$$

Finally, the spatio-temporal variance is

$$\begin{aligned} \sigma_{ST}^2 &= E_x[\text{Var}(F|x)] - \sigma_T^2 \\ &= \frac{1}{2}\Big(\text{Var}(F|x=1) + \text{Var}(F|x=2)\Big) - \sigma_T^2 \\ &= \frac{1}{2}(0 + \sigma_r^2) - \sigma_T^2 \\ &= \frac{\sigma_r^2}{2} - \frac{\sigma_r^2}{4} \\ &= \frac{\sigma_r^2}{4}. \end{aligned} \tag{S33}$$

On a relative scale, the contribution of spatio-temporal variance to total variance on the landscape can be written as

$$\frac{\sigma_{ST}^2}{\sigma_T^2 + \sigma_S^2 + \sigma_{ST}^2} = \frac{1}{2 + (-\ln(1-d) - \bar{r})^2/\sigma_r^2}. \tag{S34}$$

| | Variance Component | | |
|---|---|---|---|
| | $\sigma_T^2$ | $\sigma_S^2$ | $\sigma_{ST}^2$ |
| Expression | $\sigma_r^2/4$ | $(-\ln(1-d)-\bar{r})^2/4$ | $\sigma_r^2/4$ |

Table S1: Components of the variance in growth rates in the source sink model. $\sigma_r^2 = \mathrm{Var}(r(t))$ is the variance of the growth rates in the sink, and $d$ is the fraction of individuals that disperse from the source to the sink in a unit of time.

This shows that the absolute measure of spatio-temporal variance depends on the variance in sink growth rates, $\sigma_r^2$, whereas the contribution to the total variance on the landscape also relies on dispersal from the source, $d$, as well as the temporal average growth rate in the sink, $\bar{r}$.

#### S4 Derivation of the measure of the inflationary effect

To derive the measure of the inflationary effect, we need the growth rate of model (3) provided in the main text at the scale of the metapopulation. At the scale of the metapopulation, we can define $\bar{N} = \frac{1}{n}\sum_{i=1}^{n} N_i$ as the metapopulation density (assuming all patches are the same size). This density represents the dynamics across all space on average. The growth rate at the metapopulation scale is then

$$\frac{d\bar{N}}{dt} = \frac{1}{n}\sum_{i=1}^{n}\frac{dN_i}{dt}, \tag{S35}$$

and so is the average of the local patch growth rates. Thus, the per-capita growth rate at the metapopulation scale is

$$\frac{1}{\bar{N}}\frac{d\bar{N}}{dt} = \frac{1}{n}\sum_{i=1}^{n}\frac{1}{\bar{N}}\frac{dN_i}{dt}. \tag{S36}$$

Plugging in the expression for the patch-specific growth rates (eqn 3) of the main text into (S36) yields

$$\frac{1}{\bar{N}}\frac{d\bar{N}}{dt} = \frac{1}{n}\sum_{i=1}^{n} r_i(t)\nu_i(t) + \frac{1}{n}\sum_{i=1}^{n}\sum_{j\neq i} m_{ij}\nu_j(t) - \frac{1}{n}\sum_{i=1}^{n}\sum_{j\neq i} m_{ji}\nu_i(t), \tag{S37}$$

where $\nu_i(t) = N_i(t)/\bar{N}$ is the *relative density in patch i at time t*. The second summation on the right hand side of (S37) is the average rate of immigration across all $n$ patches (in per-capita terms). The third summation is the average rate of emigration across all $n$ patches (again, in per-capita terms). As no individuals die during dispersal in the model, the total rates of emigration and immigration are the same, i.e.,

$$\frac{1}{n}\sum_{i=1}^{n}\sum_{j\neq i} m_{ij}\nu_j(t) = \frac{1}{n}\sum_{i=1}^{n}\sum_{j\neq i} m_{ji}\nu_i(t). \tag{S38}$$

Using (S38) in (S37) yields the following expression for the per-capita growth rate at the regional scale:

$$\frac{1}{\bar{N}}\frac{d\bar{N}}{dt} = \frac{1}{n}\sum_{i=1}^{n} r_i(t)\nu_i(t) = \bar{r}(t) + \text{cov}_i(r_i(t), \nu_i(t)), \tag{S39}$$

where $\bar{r}(t) = (1/n)\sum_{i=1}^{n} r_i(t)$ is the spatial average growth rate at time $t$, and the last equality follows because the average of a product is the product of the averages plus a covariance between the two (the average of $\nu_i(t)$ is $\bar{\nu}(t) = 1$). Note that the covariance here is defined for a finite set of patches, rather than the typical interpretation in probability as a covariance over a probability measure. However, its properties should generally be identical (within sampling error) to the probabilistic interpretation when the finite set of locations under question is a random sample of space.

The metapopulation scale growth rate applies at a single point in time. The long-term growth rate in this environment is the average across the set of environmental conditions that are experienced (including their possible temporal structure). We use the expectation operator from probability theory to represent the possible conditions. Given some time $t$ far into the future, the long-term average metapopulation growth rate is

$$E_t\left[\frac{1}{\bar{N}}\frac{d\bar{N}}{dt}\right] = E_t[\bar{r}(t)] + E_t\left[\text{cov}_i\Big(r_i(t), \nu_i(t)\Big)\right], \tag{S40}$$

which gives the appropriate per-capita measure of population growth in models with continuously variable population densities in spatially and temporally varying environments.

When applied to the per-capita growth rate, the inflationary effect is the amount to which the growth rate is enhanced in reference to a hypothetical scenario without spatio-temporal variability. A suitable reference scenario is one where temporal conditions are at their average for all time. That is, each location has growth rate $\tilde{r}_i = E_t[r_i(t)]$ for all time. In this scenario, there is no spatio-temporal variation, yet there is spatial variation alone and the effects of temporal variation alone are encapsulated in $\tilde{r}_i = E[r_i(t)]$. With constant growth rates in space, the system can be described by a system of linear ordinary differential equations as

$$\frac{d\mathbf{N}}{dt} = \mathbf{A}\mathbf{N}, \tag{S41}$$

where $\mathbf{N} = (N_1, N_2, \ldots, N_n)^T$ is a column vector of local population densities, and $\mathbf{A} = (A_{ij})$ gives the rates of change in patch $i$ contributed by patch $j$. The diagonal elements, $a_{ii} = \tilde{r}_i - \sum_j m_{ji}$, describe the emigration discounted growth rate of a patch, and the off-diagonal elements, $a_{ij} = m_{ij}$ describe movement rates between patches. This model is a linear model that grows exponentially at the rate given by the dominant eigenvalue of $\mathbf{A}$ and has a stable spatial distribution given by the right eigenvector of $\mathbf{A}$.

The stable spatial distribution can be rescaled to relative density, which we label with a tilde as $\tilde{\nu}_i$ to indicate that the relative density applies to the stable distribution defined by constant growth rates $\tilde{r}_i$.

The dominant eigenvalue of **A**, which is the metapopulation scale per-capita growth rate, can be written as

$$\frac{1}{\bar{N}}\frac{d\bar{N}}{dt} = \frac{1}{n}\sum_{i=1}^{n}\tilde{r}_i\tilde{v}_i = \frac{1}{n}\sum_{i=1}^{n}\tilde{r}_i + \mathrm{cov}_i\left(\tilde{r}_i, \tilde{v}_i\right). \tag{S42}$$

In (S42), the covariance is a constant, rather than a time-varying function.

The inflationary effect is simply how much the growth rate in the actual model deviates from the reference model. Subtracting (S42) from (S40) gives an experession for the inflationary effect, which is

$$\begin{aligned}\text{Inflationary Effect} &= E_t[\bar{r}(t)] - \frac{1}{n}\sum_{i=1}^{n}\tilde{r}_i + E_t\left[\mathrm{cov}_i\Big(r_i(t), \nu_i(t)\Big)\right] - \mathrm{cov}_i\left(\tilde{r}_i, \nu_i\right) \\ &= E_t\left[\mathrm{cov}_i\Big(r_i(t), \nu_i(t)\Big)\right] - \mathrm{cov}_i\Big(E_t[r_i(t)], \tilde{v}_i\Big),\end{aligned} \tag{S43}$$

where the final line follows because

$$E_t[\bar{r}(t)] = E_t\left[\frac{1}{n}\sum_{i=1}^{n}r_i(t)\right] = \frac{1}{n}\sum_{i=1}^{n}E_t[r_i(t)] = \frac{1}{n}\sum_{i=1}^{n}\tilde{r}_i, \tag{S44}$$

and so the first two terms in the first line cancel out. The last line of equation (S43) is equation (4) of the main text.

#### S5 Details of the model of disease transmission

To illustrate the effects of spatio-temporal heterogeneity on the spread of infectious disease, we use the following two-patch SIR model:

$$\begin{aligned}\frac{dS_i}{dt} &= -\beta_i(t)\frac{S_iI_i}{N_i} - mS_i + mS_j \\ \frac{dI_i}{dt} &= \beta_i(t)\frac{S_iI_i}{N_i} - \gamma I_i - mI_i + mI_j \quad (i = 1, 2) \\ \frac{dR_i}{dt} &= \gamma I_i,\end{aligned} \tag{S45}$$

where $S_i$, $I_i$, and $R_i$ are the number of susceptible, infectious, and recovered individuals in patch $i$, respectively, $\beta_i(t)$ is the time and location-specific transmission rate, $m$ is the movement rate, and $\gamma$ is the rate of recovery from infection. In this simple form of the of model, we assume that individuals may only be in one of these three classes such that $N_i(t) = S_i(t) + I_i(t) + R_i(t)$.

As a simple illustration of asynchrony, we consider a scenario where the transmission rate oscillates between two values periodically according to the following equation (and shown in

Figure S3):

$$\beta_1(t) = \begin{cases} \beta_0 \text{ for } 2nT < t < (2n+1)T \\ \beta_0(1-\epsilon) \text{ otherwise} \end{cases}, \qquad \beta_2(t) = \begin{cases} \beta_0 \text{ for } 2nT - \tau < t < (2n+1)T - \tau \\ \beta_0(1-\epsilon) \text{ otherwise} \end{cases}, \tag{S46}$$

where $n = \{0, 1, 2, 3, \ldots\}$ is the set of natural numbers (zero inclusive), and $\epsilon$ is the proportional reduction in transmission from non-pharmaceutical interventions ($0 \leq \epsilon \leq 1$). As such, it is the effectiveness of any NPI.

The parameter $\tau$, is the time-shift between the change in state of the two patches. Patch two begins NPIs $\tau$ units of time prior to patch 1 and releases them $\tau$ units prior to patch 2 as well. Time shifts can be anywhere from 0, when the two patches are completely in sync, to $T$, when the patches are completely out of sync (note that because the square wave function is periodic, $\tau$ can take any real value, each of which has a mapping to the space $\tau \in [0, T]$).

The time shift is related to $\Omega$, the NPI overlap, with the following relationship

$$\Omega = 1 - \frac{\tau}{T}, \tag{S47}$$

where $\tau/T$ is the fraction of a half period where the patches are in different states. Each of the half periods are the same duration, so it is also the fraction of the entire period that the two patches are in different states. Therefore, $1 - \tau/T$ is the fraction of time the two patches are in the same state, i.e., the amount of overlap.

#### *S5.1 Spatio-temporal variance*

The fitness factor for this model is $\beta$, which changes over space and time. To calculate the three components of variability in $\beta$, we need the conditional spatial and temporal means and variances.

The temporal means and variances are based on the two values of the square-wave function, $\beta_0$ and $\beta_0(1-\epsilon)$, which are equally common on a cycle. Since they are equally common in both patches, the temporal average $\beta$ in patch $x$ is $E_t[\beta_x] = \beta_0(1 - \epsilon/2)$.

The pure spatial variance is the variance among patches in the temporal average transmission rate. The temporal average transmission rates are identical in the two patches, meaning there is no spatial variability. Hence,

$$\sigma_S^2 = \text{Var}_x(E_t[\beta]) = 0. \tag{S48}$$

The pure temporal variance is the variance over times in the average spatial transmission rate. To determine this, we partition time within a cycle into four periods, described below and demonstrated in Figure S3. The first time period, $t_1$, is the time when both patches have the fast

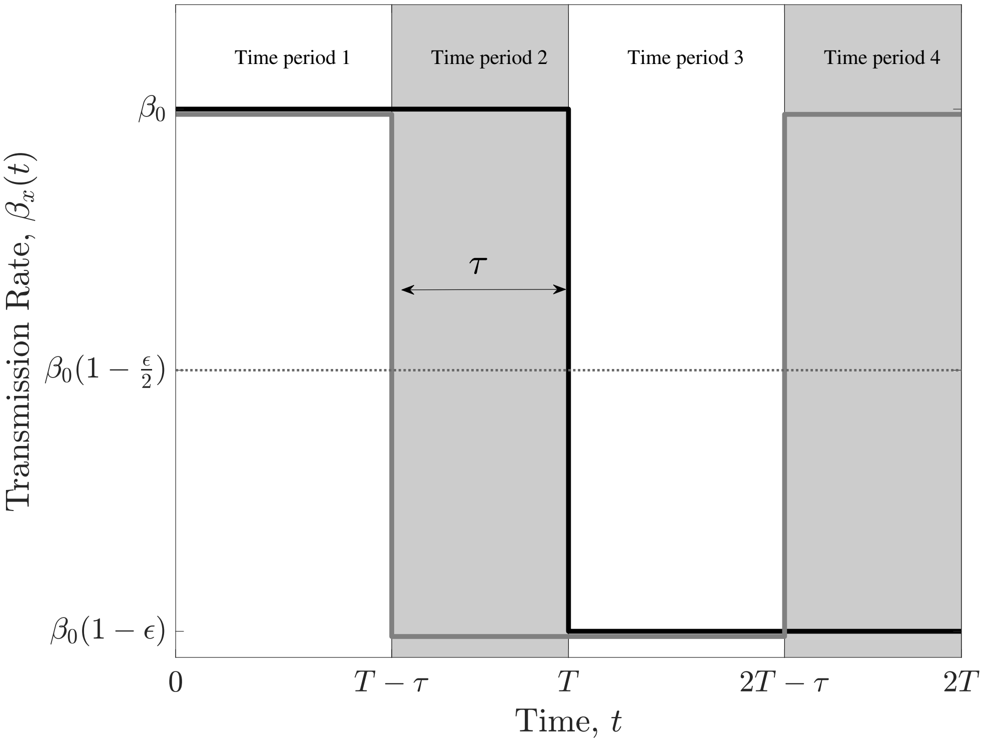


Figure S3: The square-wave function used to model differential timing of non-pharmaceutical interventions. The black line shows the transmission rate in the first patch and the gray line shows the transmission rate in the second patch. The dotted line shows the temporal average in each patch, $E_t[\beta(x,t)|x]$. Time can be partitioned into four distinct periods which fully describe the unique set of states the two patches can be in together.

transmission rate, i.e. $\beta_1 = \beta_2 = \beta_0$. The third time period, $t_3$, is the time when both patches have the slow transmission rate $\beta_1 = \beta_2 = \beta_0(1-\epsilon)$. Time periods two, $t_2$, and four, $t_4$, are those where the patches are in different states; one has transmission rate $\beta_0$ and the other has transmission rate $\beta_0(1-\epsilon)$ and the two time periods are distinguished by the identities of the patches in the different states.

Time periods one and three together comprise $\Omega$ fraction of time, and time periods two and four together comprise $1-\Omega$ fraction of time.

The average growth rate over space at any time, $E_x[\beta(x,t)|t]$ can be found by using this time partition. The average across the two patches in each time period are

$$\begin{aligned} E_x[\beta(x,t)|t \in t_1] &= \beta_0 \\ E_x[\beta(x,t)|t \in t_2] &= \beta_0\left(1-\frac{\epsilon}{2}\right) \\ E_x[\beta(x,t)|t \in t_3] &= \beta_0(1-\epsilon) \\ E_x[\beta(x,t)|t \in t_4] &= \beta_0\left(1-\frac{\epsilon}{2}\right). \end{aligned} \tag{S49}$$

The average of these spatial averages across time is

$$E_t[E_x[\beta(x,t)]] = \beta_0\left(1-\frac{\epsilon}{2}\right). \tag{S50}$$

And so the temporal variance of these spatial averages is

$$\sigma_T^2 = \mathrm{Var}_t(E_x[\beta(x,t)|t]) = \Omega\left(\frac{\beta_0\epsilon}{2}\right)^2. \tag{S51}$$

The spatio-temporal variance can be written as

$$\sigma_{ST}^2 = E_x[\mathrm{Var}_t(\beta|x)] - \mathrm{Var}_t(E_x[\beta|t]). \tag{S52}$$

To calculate this quantity, we need the spatial average of the conditional variance over time, $E_x[\mathrm{Var}_t(\beta|x)]$, as the second term on the right-hand side of the expression is the pure temporal variance.

The temporal variance in a given patch $x$ can be found by noting that the two values in a period, which are each equally common in time, deviate from the temporal average by $\pm\beta_0\epsilon/2$ . Hence, the variance over time is

$$\mathrm{Var}_t(\beta|x) = \frac{1}{2}\left(\frac{\beta_0\epsilon}{2}\right)^2 + \frac{1}{2}\left(-\frac{\beta_0\epsilon}{2}\right)^2 = \left(\frac{\beta_0\epsilon}{2}\right)^2. \tag{S53}$$

This variance is the same for both patches so that

$$E_x[\mathrm{Var}_t(\beta|x)] = \left(\frac{\beta_0\epsilon}{2}\right)^2. \tag{S54}$$

Using (S54) and (S51) in (S52) yields

$$\sigma_{ST}^2 = (1-\Omega)\left(\frac{\beta_0\epsilon}{2}\right)^2. \tag{S55}$$

Putting this on a scale of the spati-temporal variations contribution to the total variation in the transmission rate yields

$$\frac{\sigma_{ST}^2}{\sigma_S^2+\sigma_T^2+\sigma_{ST}^2} = \frac{(1-\Omega)\left(\frac{\beta_0\epsilon}{2}\right)^2}{0+\Omega\left(\frac{\beta_0\epsilon}{2}\right)^2+(1-\Omega)\left(\frac{\beta_0\epsilon}{2}\right)^2} = 1-\Omega, \tag{S56}$$

which shows that all the variation is temporal only when NPIs are exactly in sync and all the variation is spatio-temporal when NPIs are exactly out of sync.

#### *S5.2 Calculating the long-term growth rate*

We find the long-term growth rate of an invading pathogen by assuming that the fraction in the population in the infectious class is very small, which is suitable for much of the early dynamics of COVID-19 because the disease was distributed worldwide, but had infected a very small percentage of the population in any given locality of moderate spatial scale. We do this by simulating (S45) while keeping $I_i$ and $I_j$ as small constants near zero. We run this simulation over multiple cycles of $\beta_i(t)$ until a stationary distribution for $\nu_i = I_i/\bar{I}$ is reached. At that point, we simulate the dynamics of $I_i$ for more cycle and measure the long-term metapopulation rate of spread as

$$\frac{\Big(I_1(2T)+I_2(2T)\Big)-\Big(I_1(0)+I_2(0)\Big)}{2T}, \tag{S57}$$

which is the average rate of change of the infectious class at the metapopulation scale.

Figures S5.2 and S5.2 show the effects of increasing the effectiveness of NPIs and the duration of NPIs on the long-term growth rate under different overlap levels and movement scenarios.

#### *Stochastic model*

We included environmental stochasticity to this model by introducing variability to the transmission rate such that, during periods when the transmission rate does not switch (i.e., either during lockdown or during "business as usual"), the transmission follows a discretized version of the above continuous-time SIR model. The model is made stochastic by sampling $\beta_i(t)$ from eqn (S46) a truncated normal distribution with mean given by the deterministic $\beta_i(t)$ given above and variance $\sigma^2$ over one unit of time. The truncated normal is used to ensure that the transmission is not a negative value.

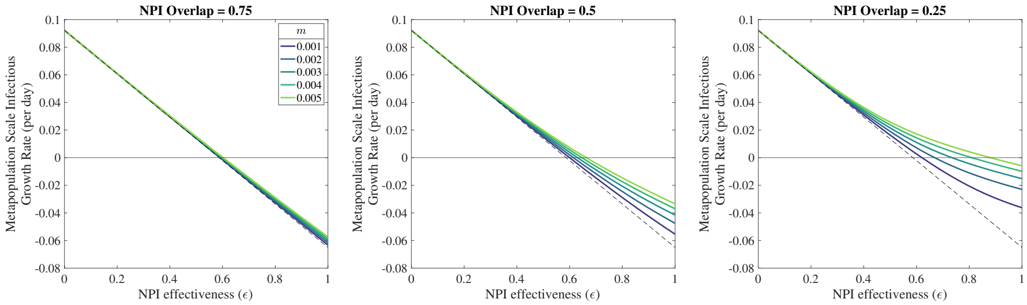


Figure S4: Metapopulation growth rate of the infectious class as a function of the effectiveness of an NPI.The dotted line shows the metapopulation spread of the disease assuming no spatio-temporal heterogeneity, which implies either coordinated NPIs or no movement. Actual spread rates lie above this line, reflecting that fact that the realized effectiveness of NPIs is diminished by asynchrony in the timing of NPIs. The difference between the dotted line any given solid line is the magnitude of the inflationary effect.

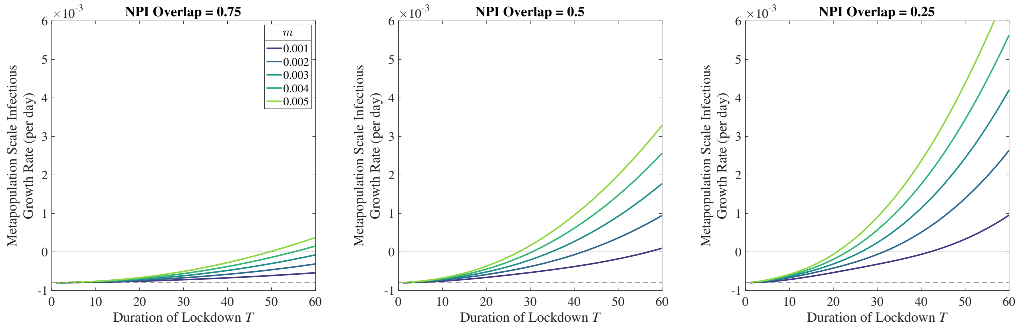


Figure S5: Metapopulation growth rate of the infectious class as a function of the duration of NPIs. Longer NPI application (and so longer durations of "business as usual") creates autocorrelated growth rates from the perspective of infectious hosts. As such, the inflationary effect is larger. The solid line gives the boundary growth rate above which the disease spreads in the metapopulation. The dashed line gives the baseline growth rate in the absence of any spatio-temporal variability in transmission.

For the stochastic model, we chose infectious duration $1/\gamma = 4.5$ days in both patches and a baseline transmission rate of $\beta_0 = 0.375$ d$^{-1}$. This corresponds to a daily growth rate (in a completely exposed population) of 1.53 d$^{-1}$, which means that cases double approximately every 4.5 days, and the disease $R_0$ value is about 1.7. We illustrate the inflationary effect with a highly effective NPI regime (95% reduction in baseline transmission) and assumed cycles of NPIs as occurring over the course of 60 days (30 days NPI regime and 30 days "business as usual").

We simulated this stochastic model by projecting the number of infectious forward with a time step of every day and chose $\sigma = \sqrt{0.01}$.
